# RFoldSearch enables fast RNA structure search beyond sequence similarity

**DOI:** 10.64898/2026.11.23.768743

**Authors:** Ruyi Chen, Wudao Yang, Hongxiang Li, Burkhard Rost, Mikael Bodén, Michael Heinzinger

## Abstract

The rapid growth of predicted RNA structure databases demands fast and sensitive structural search methods. Sequence-based approaches lose sensitivity for divergent homologues, whereas exhaustive coordinate alignment is computationally expensive. Analogous to Foldseek’s 3Di representation, we introduce RFoldSearch and 3Dr, a learned 80-state structural alphabet encoding local RNA tertiary environments. RFoldSearch combines indexed prefiltering, structure-aware token alignment and optional coordinate-based reranking. On held-out RNAsolo 2.0 queries, RFoldSearch retained 81–89% of the family-level sensitivity of exhaustive coordinate aligners and outperformed RiboSeek (RS-20) and BLASTN. On a single CPU core, RFoldSearch searched 531,888 predicted structures in 0.31 s per query, 20-fold faster than RiboSeek with its RS-20 alphabet while using 7.7-fold less peak memory. Searches of predicted-structure databases demonstrated its ability to recover structurally similar RNAs with limited detectable sequence similarity, complementing profile-based sequence search. Recurrent 3Dr words captured conserved local RNA geometries, including fragments corresponding to known RNA 3D motifs. RFoldSearch is available as open-source software, a web server and a REST API.

## 1 Introduction

RNA function is shaped by three-dimensional structure, which underlies regulation, catalysis, cellular organisation and therapeutic activity [1–3]. As experimentally determined and predicted RNA structures rapidly accumulate [4–11], identifying structural relatives is increasingly becoming a database-scale problem.

Existing approaches face a speed–sensitivity trade-off. Coordinate-based methods such as US-align [12], RMalign [13] and GTalign [14] retain high structural sensitivity, but exhaustive database searches with these methods remain computationally expensive. Sequence- and profile-based methods such as BLASTN [15] and nhmmer [16] are far faster, but lose sensitivity as structurally conserved RNAs diverge in sequence.

Discrete structural alphabets offer an alternative by converting three-dimensional structures into sequences that can be searched efficiently. Building on fast sequence-search strategies established by MMseqs2 [17], Foldseek [18] introduced the 3Di structural alphabet for rapid protein structure retrieval. Extending this approach to RNA requires representations tailored to its tertiary architecture, which is shaped primarily by base pairing, stacking and long-range interactions rather than by protein-like backbone packing [19]. Previous RNA structural alphabets have supported substructure search [20, 21], while Wang et al. [22] developed RiboSeek, which uses spatial-neighbour structural alphabets for RNA family discrimination and tertiary-structure comparison. Park et al. [23] introduced (similarly named) Riboseek for sensitive remote nucleotide sequence search. Learned RNA structure tokenisers, such as RiboSphere [24], have also been developed for representation learning and generative modelling. However, achieving sensitive and scalable RNA tertiary-structure retrieval across sequence divergence and large predicted-structure databases remains challenging.

Inspired by the success of Foldseek, and to meet the above challenges, we introduce RFoldSearch, an RNA tertiary-structure search method based on a learned 80-state structural alphabet termed 3Dr (three-dimensional RNA), which encodes local tertiary environments without nucleotide identity. RFoldSearch combines indexed token retrieval with geometric scoring and optional coordinate-based reranking (RFoldSearch-TM). On held-out experimental structures, RFoldSearch retains 81– 89% of the family-level sensitivity of exhaustive coordinate aligners and outperforms RiboSeek (RS-20) and BLASTN. Its sensitivity persists at near-background sequence identity and after excluding homologues detectable by BLASTN or nhmmer. Learned quantisation achieves retrieval accuracy comparable to *k*-means discretisation while substantially improving index selectivity, enabling sub-second search across more than half a million predicted RNA structures. On predicted structures, RFoldSearch complements sequence-based methods by retrieving sequence-divergent structural neighbours, with additional sensitivity provided by RFoldSearch-TM. Recurrent 3Dr token words further capture conserved local RNA geometry, and selected words correspond to geometrically conserved fragments of known RNA 3D motifs. These results establish RFoldSearch as a scalable approach to RNA tertiary-structure retrieval using an interpretable structural representation.

## 2 Results

### 2.1 RFoldSearch overview

RFoldSearch adapts Foldseek’s structural-alphabet search strategy [18] to RNA tertiary structures through a learned RNA-specific representation, structure-based scoring and a separately calibrated statistical model (Fig. 1A). Unlike Foldseek’s 3Di representation of protein environments, RFoldSearch uses 3Dr, an 80-state alphabet learned from local RNA tertiary geometry without nucleotide identity. Each nucleotide is represented by a 17-dimensional descriptor comprising the rigid-body pose of its nearest nonlocal spatial neighbour, g_nm_^(1)^, and a scalar context, **g**_ctx_, describing sequence separation, base-pair orientation and local contact density (see “Structural descriptors” subsection). A VQ-VAE [25] maps these descriptors to discrete 3Dr tokens (Fig. 1B).

**Fig. 1:**
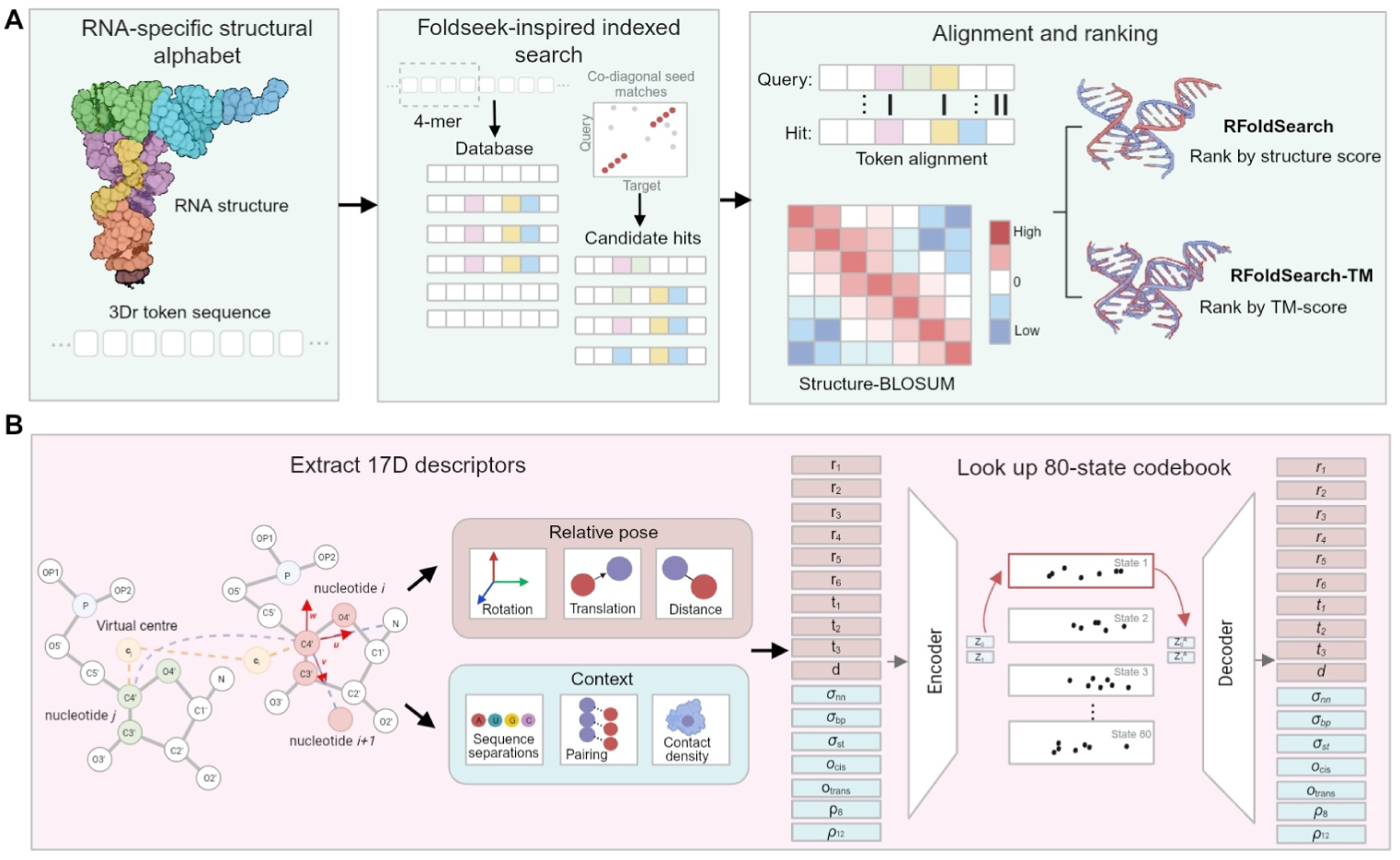
Overview of RFoldSearch. **(A)** Search pipeline. RNA tertiary structures are encoded as sequences of 3Dr structural tokens. In the schematic, matching colours denote identical 3Dr states in the query and database sequences. An independently implemented inverted-index prefilter, inspired by Foldseek’s indexed-search strategy, retrieves candidate structures using exact 4-mer matches and 4-mers containing a single token substitution. Candidates supported by at least two seeds on a consistent alignment diagonal are retained and subsequently aligned by affine-gap Smith–Waterman over the 3Dr token sequences. Token alignments are scored using Structure-BLOSUM, a substitution matrix developed here specifically for the 3Dr alphabet. In the default RFoldSearch mode, hits are ranked by the structure score *R*_str_ = *S*_SW_ TM*_a_*, which combines the Structure-BLOSUM token-alignment score with the geometric similarity of the aligned C4^′^ atom pairs. RFoldSearch-TM instead realigns retained candidates with US-align and ranks them by symmetric US-align TM-score. **(B)** 3Dr tokenisation. For each nucleotide *i*, a 17-dimensional descriptor captures the relative geometry of its nearest nonlocal spatial neighbour *j* and the local interaction context. The geometric representation uses nucleotide-local coordinate frames and base-anchored virtual centres, with arrows indicating the local reference axes. The descriptor is encoded by a vector-quantised variational autoencoder (VQ-VAE), which assigns each local environment to one of 80 learned structural states. The decoder reconstructs the input descriptor from the quantised representation.

RFoldSearch follows the general two-stage search strategy of indexed seed retrieval followed by token alignment of Foldseek, with scoring adapted to RNA structural representations. An inverted-index 4-mer prefilter considers exact and Structure-BLOSUM single-substitution neighbours, retaining candidates supported by at least two co-diagonal seeds. Retrieved candidates are aligned using affine-gap Smith– Waterman with Structure-BLOSUM, a log-odds substitution matrix estimated from aligned 3Dr token pairs in structurally similar training RNAs following the BLOSUM framework [26]. Unlike Foldseek’s combined sequence–structure scoring, RFoldSearch performs token alignment without nucleotide-identity scores to prioritise structural similarity independently of sequence conservation. The resulting token-alignment score *S*_SW_ is combined with TM*_a_*, calculated by superposing the aligned C4^′^ correspondences and averaging the query- and target-normalised TM-scores. Hits are ranked by *R*_str_ = *S*_SW_ TM*_a_*. RFoldSearch-TM optionally realigns retained candidates with US-align and ranks them by symmetric TM-score, trading additional computation for greater structural sensitivity.

Model and search parameters were selected by cross-validation using the training and validation partitions, with the test split held out (Figs. S1–S8). The codebook size was set to *K* = 80, within the range of 60–100 states where clan-level sensitivity plateaued and family-level sensitivity varied little (Fig. S3). Representative local geometries of the learned states are shown in Fig. S10.

RFoldSearch calibrates token-alignment E-values separately for experimental and predicted structure databases using target-aware Gumbel modelling, US-align-based background filtering and empirical monotone calibration. For predicted structures, RFoldSearch additionally applies composition-bias correction to the token-alignment scores (“E-values”; Fig. S15). The resulting E-values estimate the expected number of unrelated target clusters attaining at least the observed alignment score. RFoldSearch-TM ranks candidates by US-align TM-score and does not report token-alignment E-values.

### 2.2 RFoldSearch achieves sensitive RNA structural retrieval

We evaluated retrieval on held-out RNAsolo 2.0 test structures using an all-versus-database protocol, with sensitivity up to the first false positive (Sens@1FP) as the primary metric. Performance was assessed using Rfam family and clan annotations and two complementary structure-defined ground truths based on symmetric US-align TM-score and RMalign RMscore (“Methods – Evaluation on the RNAsolo benchmark”).

For annotation-defined retrieval, RFoldSearch achieved a family-level Sens@1FP of 0.655, retaining 81–89% of the sensitivity of exhaustive coordinate aligners (Fig. 2A). It outperformed RiboSeek (RS-20) and BLASTN, with no significant difference from RiboSeek (RS-80), which additionally scores nucleotide identity, or from Infernal. Coordinate-based reranking increased sensitivity to 0.687 for RFoldSearch-TM. At the clan level, RFoldSearch achieved a Sens@1FP of 0.295, outperforming Infernal, RiboSeek (RS-20) and BLASTN (Fig. 2B). Differences from exhaustive coordinate aligners were not significant, although the corresponding confidence intervals were wide because only eight clan blocks were available for statistical comparison.

**Fig. 2:**
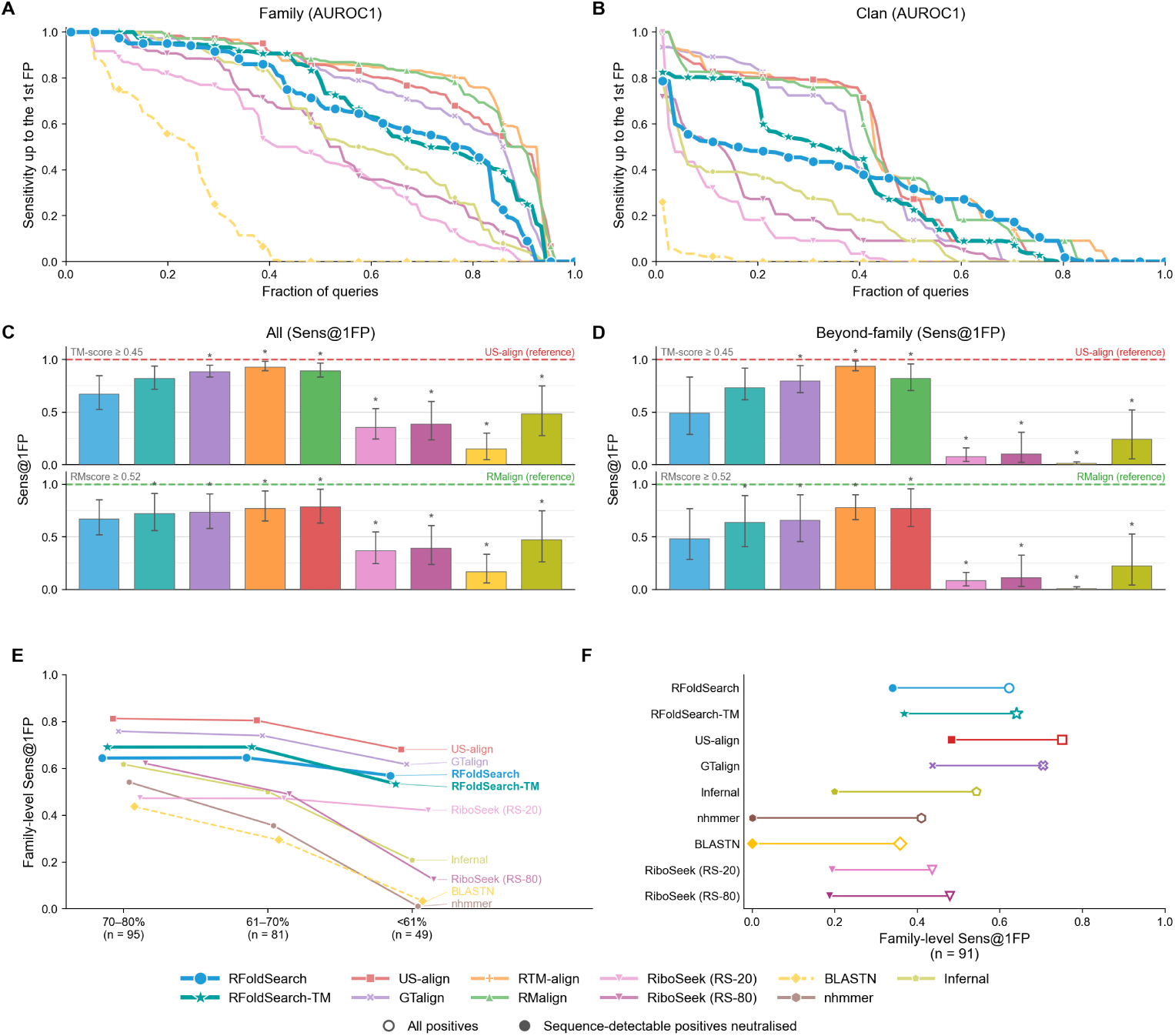
RFoldSearch retrieval performance on the held-out RNAsolo 2.0 benchmark. (A,B) Annotation-defined retrieval at the (A) Rfam family and (B) clan levels. Queries are sorted by Sens@1FP, and the area under each curve (AUROC1) equals the mean Sens@1FP. **(C,D)** Structure-defined retrieval of (C) all structural neighbours and (D) beyond-family neighbours, with positives defined by symmetric US-align TM-score 0.45 or RMalign RMscore 0.52. Error bars indicate 95% block-bootstrap confidence intervals, and asterisks mark significant differences from RFoldSearch. Dashed lines indicate the reference aligners used to define structural similarity. **(E)** Family-level Sens@1FP across sequence-identity ranges, with boundaries corresponding to the 99th (70%) and 50th (61%) percentiles of the identity distribution between unrelated RNA chains. **(F)** Family-level Sens@1FP for 91 queries before and after neutralising same-family positives detected by BLASTN or nhmmer at *E* ≤ 10.

We next used structural similarity rather than Rfam annotations to define positives. We evaluated both all structural neighbours and beyond-family neighbours, defined as structurally similar targets outside the query’s Rfam family. At a symmetric US-align TM-score threshold of 0.45, RFoldSearch achieved Sens@1FP values of approximately 0.67 for all structural neighbours and 0.49 for beyond-family neighbours (Figs. 2C and D). RFoldSearch-TM increased these values to above 0.80 and 0.70, respectively. RFoldSearch-TM reranks prefiltered candidates with US-align, the same tool that defines the ground truth. These results therefore measure how well the RFoldSearch prefilter retains US-align-defined neighbours, rather than offering a fully independent comparison with US-align. In practice, the benefit is that US-align is applied only to the small candidate set retained by RFoldSearch, not exhaustively to the entire database.

To assess dependence on the structural similarity measure, we repeated the evaluation using a quantile-matched RMalign RMscore threshold of 0.52. RFoldSearch achieved similar sensitivities of approximately 0.67 and 0.48 for all and beyond-family neighbours, respectively. It remained below exhaustive coordinate aligners while outperforming both RiboSeek alphabets, BLASTN and Infernal. Beyond family boundaries, RiboSeek (RS-80) consistently exceeded RiboSeek (RS-20), suggesting a contribution from nucleotide identity. Similar trends were observed at alternative structural-similarity thresholds (Fig. S11).

Additional ranking metrics confirmed these trends (Fig. S12). RFoldSearch achieved a family-level Recall@1 of 0.896, comparable to most methods. Its family-level mean average precision (MAP) and area under the receiver operating characteristic curve (AUROC) were higher than those of both RiboSeek alphabets, although they remained below those of US-align, RTM-align and RMalign.

Component-replacement experiments (Fig. S16) showed that the search procedure contributed most to sensitivity under structure-defined ground truth, especially for beyond-family neighbours. Its all-neighbour gain was matched by the learned substitution matrix only in the reverse order. Under Rfam annotations, the 3Dr alphabet improved family-level sensitivity in the forward order but reduced it in the reverse order, which points to an interaction with substitution scoring. Structure-based ranking added a smaller improvement.

Learned quantisation also improved index selectivity. Replacing the VQ-VAE with *k*-means clustering yielded comparable retrieval accuracy but substantially increased the number of index postings processed per query (Fig. S14). At *K* = 20, *k*-means generated 579,706 postings per query, compared with 12,598 for the VQ-VAE. Thus, the learned quantiser reduced postings by approximately 46-fold without reducing retrieval accuracy, substantially lowering the number of index entries that must be traversed and therefore the computational and memory-access cost of the prefilter.

Retrieval performance remained stable under increasingly stringent training–test partitions (Fig. S25; see Supplementary Methods). Family-level Sens@1FP was 0.595, 0.587 and 0.596 for PDB-disjoint, family-disjoint and homology-disjoint folds, respectively. Differences between the PDB-disjoint and family-disjoint evaluations were small and not significant, with similarly little change at the clan level.

### 2.3 RFoldSearch maintains retrieval sensitivity across sequence identity levels

We next examined how sequence divergence affects RFoldSearch retrieval sensitivity using the same held-out RNAsolo 2.0 benchmark. Same-family query–target pairs were stratified into three sequence-identity ranges (70–80%, 61–70% and *<* 61%), with boundaries derived from the background identity distribution of RNA pairs from different Rfam families and clans (Fig. S18A). Within each range, same-family targets outside the specified identity interval were treated as neutral (see “Retrieval performance across sequence identity levels”). The three ranges comprised 95, 81 and 49 assessable queries from 20, 15 and 8 families, respectively. Retrieval sensitivity varied substantially across methods as sequence identity decreased. While sensitivity declined sharply for nhmmer, BLASTN and RiboSeek (RS-80), RFoldSearch showed a smaller decrease, from 0.644 to 0.569. At sequence identities below 61%, RFoldSearch retained 84% of the sensitivity achieved by exhaustive US-align (Fig. 2E).

To assess retrieval beyond the detection range of conventional sequence-search methods, we neutralised same-family targets detected by BLASTN or nhmmer at *E* 10. This permissive threshold was used to exclude even weak sequence-search matches rather than to establish statistically significant homology. RFoldSearch achieved a Sens@1FP of 0.340, compared with 0.368 for RFoldSearch-TM, 0.483 and 0.437 for exhaustive US-align and GTalign, 0.199 for Infernal, and 0.194 and 0.188 for RiboSeek (RS-20) and RiboSeek (RS-80), respectively (Fig. 2F). Negative controls based on token composition, RNA length and shuffled query tokens showed substantially lower sensitivities (Fig. S18B).

At the clan level, sequence-based filtering reduced RFoldSearch Sens@1FP from 0.295 to 0.143, compared with 0.167 to 0.030 for Infernal and 0.154 to 0.046 for RiboSeek (RS-80). These changes were largely driven by the LSU and SSU rRNA clans, suggesting that clan-level performance depends strongly on the RNA groups represented in the benchmark (Figs. S18C and D).

These results demonstrate that RFoldSearch retains substantial sensitivity to remote RNA homologues, including those missed by permissive sequence-search criteria.

### 2.4 RFoldSearch alignments capture biologically meaningful nucleotide correspondences

Beyond retrieving related RNA structures, RFoldSearch also produces nucleotide-level alignments. To assess their accuracy, we compared RFoldSearch alignments against Rfam 3D seed alignments. After excluding structures used for training and validation and applying mapping and coverage filters, the benchmark comprised 313 within-family structure pairs from 24 Rfam families. Given the high sequence identity among these pairs, this benchmark primarily assesses alignment accuracy between closely related RNAs.

Under exact nucleotide-pair matching, RFoldSearch achieved a family-macro *F*_1_ of 0.790, compared with 0.734 for RiboSeek (RS-20), 0.801 for US-align and 0.944 for global sequence alignment (Fig. S20A). RFoldSearch did not differ significantly from US-align or RiboSeek but was significantly less accurate than sequence alignment (Fig. S20B).

With a positional tolerance of one nucleotide, *F*_1_ increased to 0.932 for RFold-Search, 0.876 for RiboSeek, 0.882 for US-align and 0.972 for sequence alignment. RFoldSearch then exceeded US-align by 0.050, indicating that many of its exact-match discrepancies arise from small positional offsets. The higher accuracy of sequence alignment is consistent with the sequence-based construction of Rfam reference alignments and the high sequence identity of the benchmark pairs.

Alignment accuracy varied across structure pairs, and 3Dr identity spanned a wider range than nucleotide identity (Fig. S20C–E). Representative RF00015 examples show correctly recovered correspondences alongside shifted helix or loop boundaries and alternative local alignments (Fig. S21). Together, these results show that RFoldSearch recovers Rfam-supported nucleotide correspondences, although its accuracy for remote homologues remains to be tested.

### 2.5 RFoldSearch scales to large predicted RNA structure databases

Efficient structural retrieval is essential for searching rapidly expanding RNA structure databases. To evaluate scalability, we benchmarked RFoldSearch against existing methods on two predicted-structure collections: 83,156 RNAcentral structures predicted with RhoFold+ [9] and the 531,888-structure NuFold RNA Structure Database [27] (Fig. S19). We searched nested subsets of each database using a fixed query set (see “Evaluation of runtime scalability” and “Runtime summary statistics” for details).

On RhoFold+, the mean query time of RFoldSearch increased from 0.075 s at 1,000 targets to 0.59 s at full scale (Fig. 3A). At full scale, RFoldSearch was 1,835–2,521 times faster than CPU-based coordinate aligners, 83 times faster than Infernal, 7.3 times faster than GPU-based GTalign-fast and 12 times faster than RiboSeek (RS-20), its structure-only configuration. RiboSeek (RS-80) and BLASTN were faster than RFoldSearch on this database. Coordinate-based reranking increased the mean query time of RFoldSearch-TM to 17.3 s, longer than that of GTalign. On the larger NuFold database, RFoldSearch required 0.31 s per query on a single CPU core, making it 20, 43 and 1.5 times faster than RiboSeek (RS-20), GTalign-fast and BLASTN, respectively, whereas RiboSeek (RS-80) required 0.086 s per query (Fig. 3B). RFoldSearch-TM was also faster than GTalign-fast on this database.

**Fig. 3:**
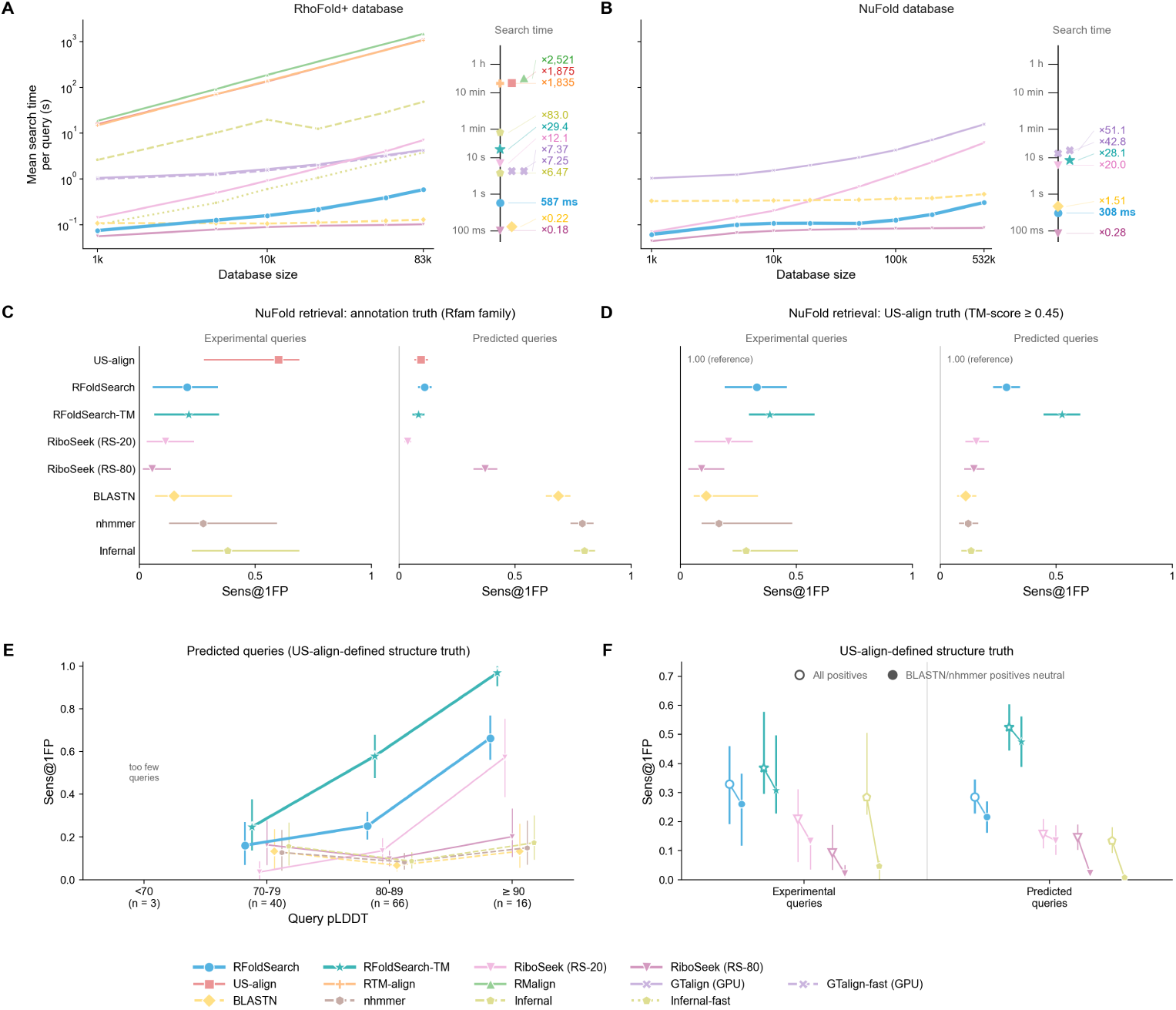
Search speed and retrieval on predicted-structure databases. (A,B) Mean search time per query against database size for the RhoFold+ and NuFold databases, respectively. Right: mean search time on the full database, expressed relative to RFoldSearch. **(C,D)** Sens@1FP under annotation-defined (Rfam family) and structure-defined (symmetric US-align TM-score 0.45) ground truth, respectively, for experimental and predicted queries. **(E)** Structure-defined Sens@1FP for predicted queries, stratified by mean query pLDDT. **(F)** Structure-defined Sens@1FP before and after excluding positives detected by BLASTN or nhmmer (*E* 10). Sens@1FP was evaluated with a retrieval depth capped at 200 non-neutral hits and normalised by the smaller of 200 and the total number of positives. Retrieval points show query means with 95% confidence intervals from family-level hierarchical bootstrapping.

RFoldSearch query time increased sublinearly with database size on both bench-marks (Figs. S13A–D). On NuFold, the fitted scaling exponent was 0.21 for RFold-Search and 0.72 for RiboSeek (RS-20), indicating that RFoldSearch runtime increased more slowly as the database grew. Its mean speed advantage over RS-20 increased from 1.1-fold at 1,000 structures to 20-fold at full scale (Fig. S17A). This mean advantage was not uniform across queries. At full NuFold scale, RFoldSearch was faster than RS-20 for 189 of the 300 paired queries, with a median speed-up of 2.3-fold. The larger 20-fold difference between the mean query times was driven by a subset of long queries that were particularly slow for RS-20 (Fig. S17B). In contrast, RiboSeek (RS-80) was 1.3–5.7 times faster than RFoldSearch across the evaluated database sizes.

Search time also increased with query length, measured in 3Dr tokens, with log–log slopes of 2.6 on RhoFold+ (*r*^2^ = 0.67) and 1.1 on NuFold (*r*^2^ = 0.80) (Figs. S13E and F). The slowest 10% of queries accounted for 48% and 42% of the total RFoldSearch query time on RhoFold+ and NuFold, respectively.

Peak memory of a single search process on the full databases is listed in Table S10. RFoldSearch required 5.5 GB on RhoFold+ and 12.7 GB on NuFold, as it keeps database coordinates in memory for structure scoring. RiboSeek (RS-20) required 94.6 and 97.5 GB, respectively, corresponding to 17-and 7.7-fold higher peak memory than RFoldSearch. RiboSeek (RS-80) required 1.4 and 7.0 GB, respectively.

Overall, these results show that RFoldSearch maintains subsecond mean query times when searching hundreds of thousands of predicted structures. Compared with the structure-only RiboSeek RS-20 configuration, RFoldSearch scaled more favourably and required substantially less peak memory. RiboSeek (RS-80), which incorporates nucleotide identity, remained faster and more memory-efficient than RFoldSearch. RFoldSearch’s favourable scaling likely reflects its selective *k*-mer indexing and bounded candidate alignment.

### 2.6 RFoldSearch detects structural similarities beyond sequence search in predicted RNAs

We next evaluated RFoldSearch on predicted RNA structures using a NuFold bench-mark. The database contained 33,043 structures from 47 organisms, spanning 2,264 Rfam families. It included up to 50 annotated structures per family and 15,320 unannotated structures matched by organism, length and prediction confidence (see “Evaluation on the NuFold benchmark” in Methods). Queries comprised 69 held-out experimental structures and 150 predicted structures. Retrieval was assessed against two ground truths. Annotation-defined positives shared the query’s Rfam family, with same-family targets from the query’s own organism treated as neutral for predicted queries. Structure-defined positives had a symmetric US-align TM-score 0.45 to the query, with RMscore-defined positives providing an alternative structural evaluation.

The two ground truths yielded similar method rankings for experimental queries but contrasting rankings for predicted queries (Figs. 3C and D). Under annotation-defined evaluation, sequence-based methods performed best for predicted queries.

Infernal achieved a Sens@1FP of 0.80, compared with 0.37 for RiboSeek (RS-80), which incorporates nucleotide identity, and 0.11 for RFoldSearch. Exhaustive US-align achieved only 0.09 for predicted queries, compared with 0.60 for experimental queries. Cross-organism retrieval was also less sensitive than within-organism retrieval across methods (Fig. S24A). The low sensitivity of even exhaustive US-align suggests that Rfam family membership was often not reflected in similarity between the predicted coordinates. Rfam families are defined by sequence and covariance-model signals rather than by tertiary structure, and predicted members of a family may differ in global fold because of prediction errors or within-family structural heterogeneity; sequence-detectable homologues therefore need not share strong global three-dimensional similarity. Consistently, treating BLASTN-or nhmmer-detected positives as neutral left a small and difficult residual set and reduced nearly all methods to low sensitivity, indicating that structure-based retrieval did not subsume sequence-based family recovery on this benchmark (Fig. S22).

In contrast, under structure-defined evaluation, RFoldSearch outperformed sequence-based methods for predicted queries. RFoldSearch achieved a Sens@1FP of 0.29 for predicted queries, outperforming Infernal and both RiboSeek alphabets. RFoldSearch-TM further increased sensitivity to 0.52 through US-align reranking, although this evaluation favours TM-score-based ranking because the same similarity measure defines the structural positives. These trends were consistent across evaluation metrics, RMscore-defined positives and structural similarity thresholds (Fig. S23).

Structure-defined retrieval was strongly associated with prediction confidence (Fig. 3E). For predicted queries, RFoldSearch Sens@1FP increased from 0.16 at pLDDT 70–79 to 0.66 at pLDDT 90, while RFoldSearch-TM increased from 0.25 to 0.97. Because sequence-based methods do not use predicted coordinates, their performance differences across confidence bins likely reflect differences in the underlying query sets rather than a direct dependence on structural confidence. Experimental queries, which carry no prediction confidence, depended similarly on the confidence of their targets: structural retrieval was weak for targets below pLDDT 80 but competitive with sequence search for high-confidence targets (Fig. S24B).

After treating structure-defined positives detected by BLASTN or nhmmer at *E* 10 as neutral, RFoldSearch retained Sens@1FP values of 0.22 and 0.26 for predicted and experimental queries, respectively, while RFoldSearch-TM reached 0.48 and 0.31 (Fig. 3F). Thus, RFoldSearch recovered a subset of structural neighbours beyond sequence-based retrieval. In a representative 5S rRNA example, RFoldSearch ranked a high-confidence target at position 11 that BLASTN, nhmmer and Infernal did not detect under the evaluated settings (Fig. S26).

Collectively, these results show that profile-based sequence search remains more effective for recovering Rfam family membership using nucleotide sequence alone. RFoldSearch addresses a complementary task by retrieving geometrically similar and sequence-divergent neighbours from predicted-structure databases, particularly when prediction confidence is high. Coordinate reranking further increases sensitivity under structure-defined evaluation by applying US-align to the candidates retained by the RFoldSearch prefilter.

### 2.7 Recurrent 3Dr words correspond to conserved local RNA structures

To assess whether recurrent 3Dr words capture local structural similarity, we identified 2,054 four-token words recurring across Rfam families and clans in the curated RNAsolo 2.0 set (see “Recurrent token-word detection and structural annotation”). Occurrences from different clans had lower median pairwise RMSDs than frequency-matched words, with larger differences relative to sequence-matched or random windows. This pattern was consistent across the descriptor anchors and the base-inclusive atom set (Figs. 4A and B).

**Fig. 4:**
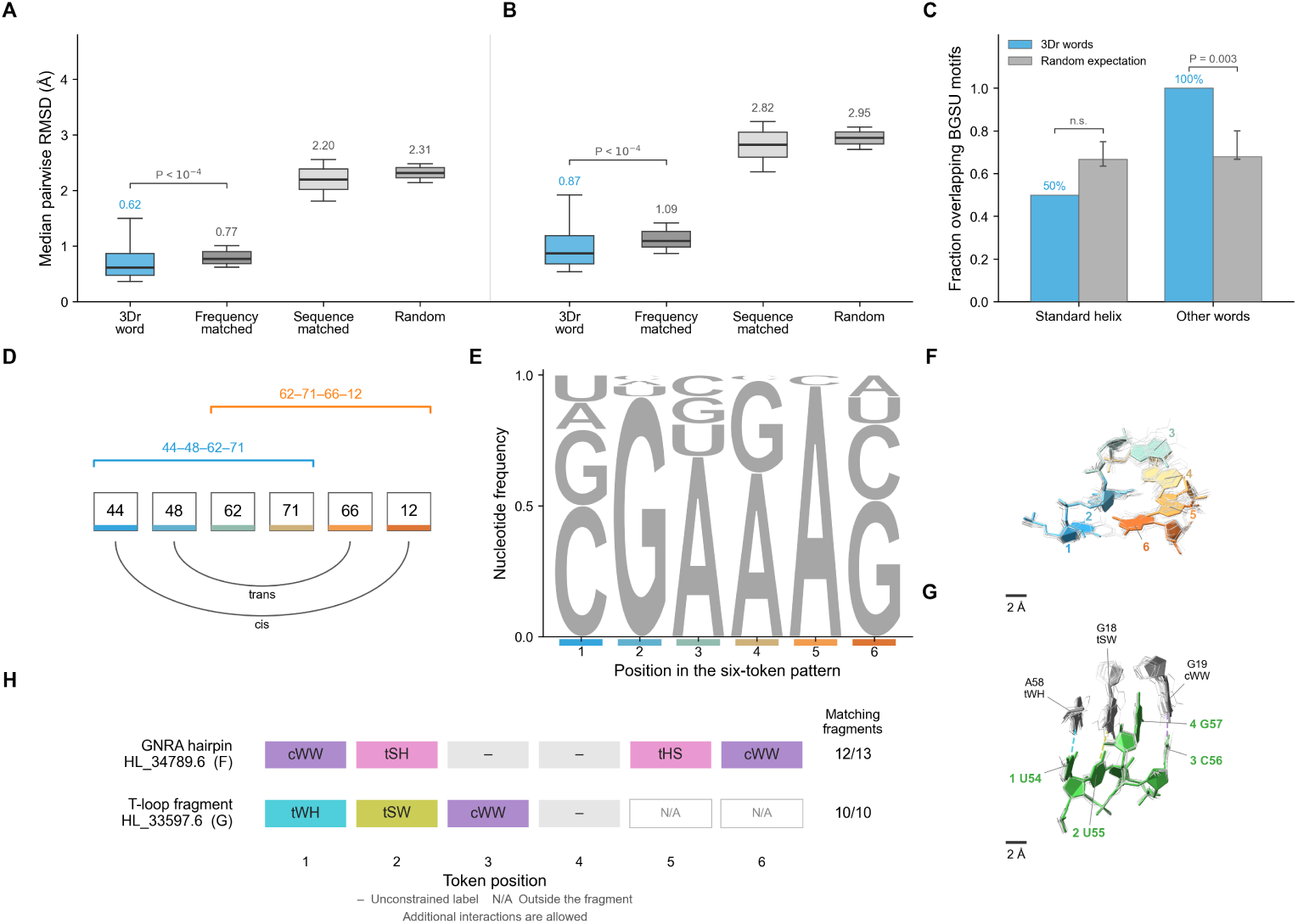
Recurrent 3Dr words correspond to conserved local RNA structures. (A,B) Per-word median cross-clan RMSDs for recurrent 3Dr words and controls, using (A) five descriptor anchors or (B) base-inclusive atoms. Boxes show interquartile ranges; lines and numbers indicate medians; whiskers span the 10th– 90th percentiles. *P* values compare recurrent and frequency-matched words using two-sided unique-word permutations within occurrence-frequency bins. **(C)** Median per-word fraction of occurrences overlapping BGSU hairpin or internal-loop motifs by at least one nucleotide, for standard-helical and other words. Grey bars show null medians; error bars span the central 95% of null replicates. *P* values are from one-sided Monte Carlo tests; n.s., not significant. **(D)** Overlapping words 44-48-62-71 and 62-71-66-12 form the six-token pattern 44-48-62-71-66-12. **(E)** Per-position nucleotide frequencies across the 48 instances of 44-48-62-71-66-12. **(F,G)** Base-inclusive superpositions of (F) 13 six-nucleotide HL 34789.6 fragments and (G) 10 four-nucleotide HL 33597.6 fragments. Representatives are coloured; other instances are thin grey sticks. In G, green residues U54–G57 and grey external partners A58, G18 and G19 use deposited numbering in the representative (PDB 4GCW, chain B). Dashed connectors denote pairing classes from focal residue to partner, not hydrogen bonds. The views share a physical scale; scale bars, 2 Å. **(H)** Required DSSR Leontis– Westhof interaction labels by token position, ordered from each focal nucleotide towards its partner. Fractions indicate the proportion of fragments containing all required labels. Dashes indicate unconstrained positions; N/A marks positions outside the four-nucleotide fragment. Additional interactions are allowed.

We next asked whether these structurally conserved words overlap annotated RNA motifs. When overlap required at least one shared nucleotide, the median proportion of occurrences overlapping BGSU hairpin or internal-loop motifs was 85.7% per word, compared with 66.7% under the occurrence-matched null. Enrichment was detected for words classified as other words, but not for standard-helical words (Fig. 4C). No enrichment was detected when at least three or all four nucleotides were required to overlap an annotated motif. The association therefore depended on the overlap criterion and did not establish that recurrent words generally cover complete motifs or their interaction cores.

Despite the limited overlap enrichment, selected overlapping words covered all aligned positions of an annotated motif. The words 44-48-62-71 and 62-71-66-12 formed the six-token pattern 44-48-62-71-66-12, observed in 48 instances from 17 Rfam families and five clans. All shared the same two annotated pair relationships: positions 1 and 6 formed a cis pair, and positions 2 and 5 formed a trans pair (Fig. 4D). Although nucleotide identity is not encoded by 3Dr, 44 instances had a GNRA sequence at positions 2–5, and 37 had G–C or C–G at the closing-pair positions 1 and 6. The sequence logo summarises nucleotide frequencies at each position (Fig. 4E). Thirteen instances covered all six aligned nucleotides of annotated HL 34789.6 hairpins and showed conserved base-inclusive geometry (Fig. 4F); 12 contained all required labels in the positional DSSR signature cWW,tSH,--,--,tHS,cWW (Fig. 4H) [28, 29]. Labels are ordered from each focal nucleotide towards its partner: cWW denotes cis Watson–Crick/Watson–Crick pairing, whereas tSH and tHS describe the trans sugar-edge/Hoogsteen pair from its two respective ends. Dashes denote positions without a required interaction label.

A separate recurrent word, 48-24-39-39, was associated with the T-loop group HL 33597.6. Of its 11 occurrences in the BGSU overlap analysis, ten were contained within annotated occurrences of this motif group, covering four of its nine aligned nucleotides. These fragments showed conserved base-inclusive geometry and contained all required labels in the positional DSSR signature tWH,tSW,cWW,--. The first three positions paired with nucleotides outside the four-token window. In the displayed representative (PDB 4GCW, chain B), the window comprises deposited residues U54– G57, and the external partners are A58, G18 and G19, respectively (Figs. 4G and H). A four-token window specifies four consecutive token positions, but each token encodes structural context that can include nucleotides outside that window.

These results show that recurrent 3Dr words capture conserved local RNA geometry, while selected words and their combinations correspond to geometrically conserved fragments of known RNA motifs.

## 3 Discussion

RFoldSearch enables scalable RNA tertiary-structure retrieval by combining a learned structural alphabet with indexed candidate generation and alignment. By retaining substantial structural sensitivity at a fraction of the computational cost of exhaustive coordinate alignment, RFoldSearch makes systematic screening of large predicted RNA structure collections more practical. This allows researchers to prioritise potentially related structures for further analysis. RFoldSearch-TM provides an intermediate option by reranking retrieved candidates through coordinate alignment, trading speed for greater structural sensitivity.

Several design choices contribute to this balance. The structure score adds geometric agreement to token alignment, reducing matches that share local motifs but differ in overall architecture. Although *k*-means achieved comparable retrieval sensitivity, it required approximately 46-fold more index postings to be accessed per query, highlighting the advantage of learned quantisation in improving index selectivity. By reducing index traversal and associated memory accesses, learned quantisation lowers the computational cost of prefiltering before candidate alignment. Structure-BLOSUM further improves retrieval by tolerating substitutions between geometrically similar structural states. These findings highlight how structure-aware scoring, selective indexing and substitution-based alignment contribute to retrieval sensitivity and efficiency.

The distinction between sequence and structure became clearest for sequence-divergent and predicted RNAs. RFoldSearch retained substantial sensitivity after sequence-detectable homologues were neutralised, whereas sequence-based methods and RiboSeek showed sharper declines. On predicted structures, profile-based sequence search remained superior for recovering Rfam family members, while RFoldSearch retrieved structurally similar targets missed by sequence search, particularly among high-confidence predictions. Coordinate reranking further increased sensitivity under structure-defined evaluation. These results highlight the complementary roles of sequence and structure search: sequence profiles recover established homology, whereas structural retrieval identifies three-dimensional similarity that may persist beyond detectable sequence similarity.

The learned alphabet also captures conserved local RNA geometry. Recurrent 3Dr words preserved local structural similarity across Rfam families and clans. Although enrichment for annotated motifs was limited to a permissive overlap criterion, selected words and their combinations corresponded to geometrically conserved fragments of GNRA hairpins and T-loops. Thus, 3Dr provides not only a representation for efficient retrieval but also a means of identifying recurrent local structural patterns.

RFoldSearch currently operates on individual RNA chains and remains sensitive to the quality of predicted structures. Furthermore, structural similarity does not necessarily imply evolutionary homology or shared biological function, particularly for predicted models. Nevertheless, the ability to search large RNA structure collections rapidly provides a practical complement to sequence-based retrieval and exhaustive coordinate alignment, with potential applications in systematic structural screening and annotation.

## Methods

### Dataset curation and redundancy reduction

We curated a non-redundant RNA structure dataset from RNAsolo 2.0 [4, 30], a cleaned PDB-derived collection of experimentally determined RNA three-dimensional structures, using the version downloaded on 22 April 2026. Each structure was represented as a single RNA chain. For each nucleotide, we extracted the C3^′^, C4^′^ and O4^′^ atom coordinates, which define the local backbone frame. Nucleotides missing any of these atoms were retained in the coordinate tensor but marked invalid in a per-position binary mask.

Redundancy was reduced in two stages, following the chain-level filtering strategy of RhoFold+ [9]. First, RNAsolo chains were intersected with the BGSU RNA 3D Hub non-redundant list [31], release version 4.38 (22 April 2026), at the 4.0 Å resolution threshold. Multi-chain Integrated Functional Element (IFE) classes were excluded because RFoldSearch models single-chain RNA structures. For each retained single-chain equivalence class, all member chains were kept, yielding 9,394 candidate RNA chains. Second, these chains were clustered with CD-HIT-EST [32] at an 80% sequence-identity threshold to remove residual chain-level redundancy beyond BGSU IFE grouping. One representative chain was selected from each sequence cluster, giving 891 non-redundant chain representatives. After excluding chains shorter than 20 nucleotides, the final dataset contained 885 representative RNA chains.

### Rfam family and clan annotation

Rfam family annotations were assigned using the Rfam pdb full region.txt cross-reference file. For chains matched to multiple Rfam families, we selected a deterministic primary family label, defined as the hit with the lowest E-value, for retrieval evaluation and reporting.

To evaluate retrieval beyond individual families, each primary Rfam family was mapped to its corresponding Rfam clan accession using the Rfam clan membership.txt file. Chains whose primary family had no clan assignment were retained in the searchable database but were not used as clan-level queries. Chains without a primary Rfam family assignment (Rfam-unassigned) were retained as neutral search targets.

### Data partitioning

Model development used a fixed PDB-disjoint train/validation/test split, with all RNA chains from each PDB entry assigned to the same partition. This prevented overlap of PDB entries but not necessarily sequence- or structure-related chains across partitions. Rfam family and clan overlap was permitted, with generalisation assessed separately using stricter family- and homology-disjoint splits (“Retrieval performance under alternative data splits”).

The split comprised 610 training, 85 validation and 190 test chains from 556, 79 and 159 PDB entries, respectively. Rfam family annotations covered 359, 43 and 114 chains (82, 17 and 35 families), while clan annotations covered 304, 36 and 95 chains (25, 7 and 11 clans) (Table S1).

### Structural descriptors

#### Local coordinate frame

We construct a nucleotide-local coordinate frame using the C3^′^, C4^′^ and O4^′^ atoms, adopting the atom selection of Anand et al. [33]. With C4^′^ as the origin, the vectors C3^′^ C4^′^ and C4^′^ O4^′^ are orthogonalised using Gram–Schmidt to define the local axes. We use ribose atoms rather than base atoms because they provide a consistent reference across canonical and many chemically modified nucleotides.

Let 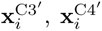 and 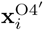*′* denote the coordinates of the C3^′^, C4^′^ and O4^′^ atoms, respectively, in nucleotide *i*. The first basis vector, **e**_1_*_,i_*, is defined along the local backbone direction as

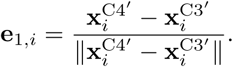

The second basis vector is obtained by projecting 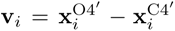 onto the plane orthogonal to **e**_1*,i*_:

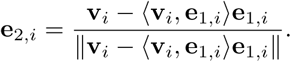

The third orthonormal basis vector is then obtained using the right-hand rule:*′*

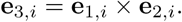

Together, the rotation matrix **R***_i_*= [**e**_1*,i*_, **e**_2*,i*_, **e**_3*,i*_] and the origin 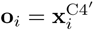 define the local rigid frame *T_i_* = (**R***_i_,* **o***_i_*) of nucleotide *i*.

#### Two descriptor families

Each nucleotide descriptor combines two complementary feature families: FRAME descriptors encode the relative rigid-body pose to a selected partner, while SCALAR descriptors summarise sequence separations, base-pairing geometry and local contact density. They capture three interaction modes that organise RNA tertiary structure: base pairing, base stacking and non-local spatial proximity.

#### FRAME descriptors

Given two nucleotides *i* and *j* with local rigid frames *T_i_* = (**R***_i_,* **o***_i_*) and *T_j_* = (**R***_j_,* **o***_j_*), the pose of *j* in the frame of *i* is given by the relative translation and rotation

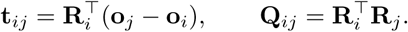

From these we obtain the inter-frame distance *d_ij_* = ||**t***_ij_||* _2_ and the unit direction ^**t***_ij_* = **t***_ij_/*||**t***_ij_||*_2_ in the frame of *i*. The first two columns of **Q***_ij_* ∈ SO(3) give the continuous 6-D rotation representation of Zhou et al. [34]. RFoldSearch stores their entries in row-major order,

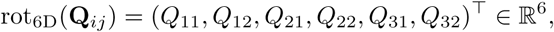

where *Q_ab_* denotes entry (*a, b*) of **Q***_ij_*. The 10-dimensional FRAME descriptor concatenates the distance, direction and rotation representation,

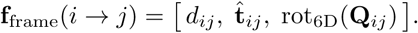

*Spatial neighbour* g_nm_^(1)^ (10-D). The first FRAME block describes the closest nucleotide in space that is not a near neighbour in sequence. Inter-nucleotide distances are measured between virtual centres **c***_i_*, which are placed near C4^′^ and shifted slightly towards the base along the direction of the glycosidic nitrogen (N9 for purines, N1 for pyrimidines). The construction is detailed in the Supplementary Methods. For nucleotide *i*, the neighbour *j* is the nucleotide whose virtual centre is closest among those at least four positions away in sequence. Applying the FRAME template to the pair (*i, j*) yields the 10-dimensional feature **g**^(1)^ R^10^. The weights defining the virtual centre were fixed in advance rather than tuned, and retrieval was insensitive to their values (Fig. S7).

#### SCALAR descriptors

The 7-dimensional scalar context descriptor **g**_ctx_ comprises signed sequence separations to the spatial-neighbour, base-pair and stacking partners, a one-hot Leontis–Westhof cis/trans indicator and two-shell non-sequential contact densities.

*Sequence separations* (*σ*_nn_*, σ*_bp_*, σ*_st_) (3-D). For each nucleotide *i*, we define three interaction partners *p* nn, bp, st , corresponding to its nearest nonlocal spatial neighbour, base-pairing partner and base-stacking partner, respectively. The nearest spatial neighbour is defined consistently with that used in the FRAME block. Base-pairing and base-stacking interactions are identified using Biotite [35]. When multiple candidate partners are identified for a given interaction type, the partner with the smallest sequence separation from nucleotide *i* is selected.

Let *j_p_*(*i*) denote the sequence index of partner *p* associated with nucleotide *i*. We define the signed sequence separation as

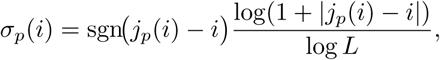

where *L* denotes the RNA sequence length. Normalisation by the maximum possible sequence separation constrains *σ_p_*(*i*) to [−1, 1]. If no partner of type *p* is identified for nucleotide *i*, we set *σ_p_*(*i*) = 0.

*Cis/trans glycosidic-bond orientation* (*o*_cis_*, o*_trans_) (2-D). For each nucleotide with an identified base-pairing partner, we encode the Leontis–Westhof glycosidic-bond orientation [29, 36], which classifies a base pair according to whether the two glycosidic bonds lie on the same (*cis*) or opposite (*trans*) sides of the inter-base axis. The orientation is determined using biotite.structure.base_pairs_glycosidic_bond [35] and represented as a two-dimensional one-hot vector,

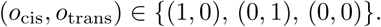

Here, (1, 0) and (0, 1) denote cis and trans orientations, respectively, whereas (0, 0) is assigned when no base-pairing partner is identified or when the glycosidic-bond orientation cannot be determined.

*Non-sequential contact density* (*ρ*_8_*, ρ*_12_) (2-D). We summarise local packing with a two-shell contact density, analogous to the coordination-number descriptors used in proteins [37]. For each nucleotide *i* and radius *r* 8, 12 Å, we count the non-sequential nucleotides whose C4^′^ reference points lie within *r*, normalised and capped at one,

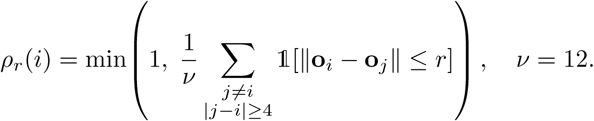

The |*j - i|* 4 restriction excludes trivial contacts arising from chain connectivity. Concatenating these three blocks gives the 7-dimensional context descriptor,

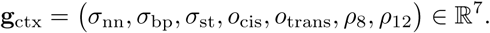

#### Descriptor assembly

The final RFoldSearch descriptor concatenates the nearest-spatial-neighbour FRAME block with the SCALAR context block,

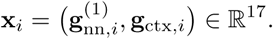

The larger 49-dimensional descriptor used for feature-ablation experiments is described in the Supplementary Methods.

### VQ-VAE learning of RNA structural tokens

We learn a discrete RNA structural alphabet using a VQ-VAE [25]. Each nucleotide is represented by the 17-D production descriptor 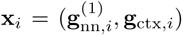. For implementation convenience, it is stored in the 49-dimensional ablation layout, with inactive dimensions zeroed and excluded from the reconstruction loss.

The encoder consists of LayerNorm followed by a feed-forward neural network (FNN) with sigmoid linear unit (SiLU) activation. It maps **x***_i_* to a latent representation **z***_i_* = *f*_enc_(**x***_i_*) ∈ R*^d^z* , with *d_z_* = 128. The encoder output is L2-normalised,

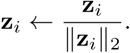

The codebook contains *K* prototype embeddings, 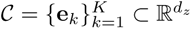, which are also L2-normalised. The structural token assigned to nucleotide *i* is the index of its nearest codebook embedding,

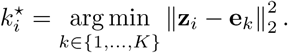

Because both **z***_i_* and **e***_k_* lie on the unit hypersphere, this assignment is equivalently given by maximum cosine similarity. The corresponding quantised latent vector is **z***_q,i_* = **e***_k_\**. A decoder FNN reconstructs the canonical descriptor, **x**^*_i_* = *f*_dec_(**z***_q,i_*).

The model was trained with a reconstruction objective defined only on the active production blocks,

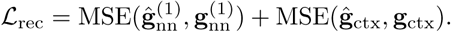

The encoder was additionally regularised by the VQ commitment loss

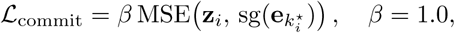

where sg( ) denotes the stop-gradient operator and *β* is the commitment-loss weight. The total training objective is

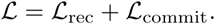

Decoder gradients were passed to the encoder using the straight-through estimator, while the L2-normalised codebook embeddings were updated by exponential moving averages (EMA) with a decay of 0.99 [38].

Descriptor normalisation statistics were estimated from the training split only and then applied to validation, test and database structures. RFoldSearch uses a two-layer encoder and a four-layer decoder, with hidden width *d_h_* = 128 (Fig. S4 and Table S5). Models were trained for 50 epochs with AdamW (learning rate 3 10^−4^, weight decay 10^−2^), cosine learning-rate decay with 200 linear warm-up steps, gradient clipping at norm 1 and a batch size of 512 nucleotide descriptors. For each random initialisation, the checkpoint with the lowest validation loss was retained. We trained 30 independently initialised models, retaining the checkpoint with the lowest validation reconstruction loss for each. Models were then evaluated on validation retrieval using the deployed search pipeline and a training-derived Structure-BLOSUM matrix. We selected the model with the highest combined family- and clan-level Sens@1FP based on Rfam annotations.

### k-mer prefiltering

Candidate targets were identified using a *k*-mer prefilter [18]. For each query 4-mer, we retrieved its exact match and one-substitution neighbours with a Structure-BLOSUM score of at least 3.0. Hits were grouped by diagonal offset, *d* = *q*_pos_ *t*_pos_, and targets were ranked by the number of hits on their best diagonal, requiring at least two matches. The retained candidates were subsequently aligned by token Smith–Waterman and ranked by structure score. Ties at the candidate boundary were resolved deterministically using a seeded hash of the chain identifier. Prefilter parameters were selected by ablation (Figs. S6 and S8).

### 3Dr substitution matrix

Candidates were aligned to the query by Smith–Waterman local alignment [39] with affine gap penalties [40] (gap opening 2, extension 0.5), using the Structure-BLOSUM matrix as substitution scores and a Numba-compiled implementation. Following the log-odds construction used by Foldseek for 3Di [18], token co-occurrences were estimated from structurally similar chain pairs rather than from Rfam labels. We selected unique unordered pairs within the model-training partition with a symmetric US-align TM-score of at least 0.5. The 3Dr sequences of each pair were then aligned by local Smith–Waterman with an identity scoring matrix and gap penalties of −11 (opening) and −1 (extension). Every non-gap traceback column, whether matched or mismatched, added one count to both *C_ij_* and *C_ji_*

From these counts we computed

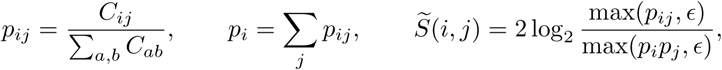

with *ɛ* = 10^−12^ to avoid log 0, and rescaled the matrix to unit mean diagonal,

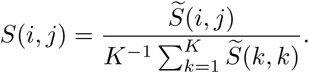

Among the tested scoring schemes, Structure-BLOSUM gave the highest mean family-level Sens@1FP, within one standard deviation of the Rfam-supervised alternative (Supplementary Figs. S5 and S9).

### Structure score

Candidates passing the *k*-mer prefilter were aligned by affine-gap token Smith– Waterman, yielding the alignment score *S*_SW_ and nucleotide correspondences. For an alignment with *L*_aln_ non-gap pairs, let *L_Q_* and *L_T_* be the numbers of valid-coordinate nucleotides in the query and target, and let *r_i_* be the post-superposition C4^′^ distance for pair *i*. We defined

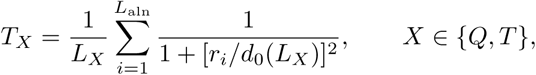

with

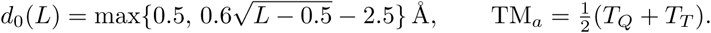

The aligned coordinates were superposed by iterative Kabsch refinement for up to five iterations, with initialization and inlier-update details provided in the Supplementary Methods. The retained value was the maximum TM*_a_* across the refinement trajectory. Alignments with fewer than three nucleotide pairs were assigned TM*_a_* = 0. Hits were ranked by

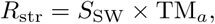

combining token-level similarity with spatial consistency.

### Predicted structure databases

Two predicted RNA structure databases were used for large-scale runtime evaluation. The NuFold RNA Structure Database [27] (version 1-13-2026) comprises RNAcentral sequences of up to 400 nt, with structures predicted using the MSA-based NuFold [10] model. We retained the rank-1 model for each entry and applied the study-wide minimum length threshold of 20 nt, yielding 531,888 structures.

We additionally constructed a RhoFold+ [9] database from RNAcentral [41]. Rfam annotations [42] were used only for family-stratified sampling. Canonical RNA sequences of 20–500 nt were retained, with up to 30 sequences sampled per Rfam family, yielding 83,156 sequences from 4,026 families. Structures were predicted with RhoFold+ in single-sequence mode using single-precision inference on an NVIDIA RTX 4090 GPU, without Amber relaxation.

### Evaluation on the RNAsolo benchmark

Retrieval was evaluated on a held-out RNAsolo 2.0 benchmark using Foldseek’s allversus-database protocol. Each chain was queried against all other chains, with self-hits excluded. The search database contained 885 RNA chains, including 516 with Rfam annotations and 369 without. Rfam-unassigned chains were retained to preserve the full benchmark database and allow their evaluation under structure-defined ground truths, but were treated as neutral in annotation-defined evaluation.

RFoldSearch used the learned 80-state 3Dr alphabet, a 4-mer prefilter and affine-gap token Smith–Waterman alignment of the top 200 candidates, followed by structure-score ranking. Results were averaged over ten seeded tie-orderings. RFoldSearch-TM reranked the same candidates using symmetric US-align TM-score. Coordinate-based comparators included US-align, RMalign, RTM-align and GTalign, ranked by their respective symmetric structural scores. RiboSeek used its RS-80 (structure × nucleotide identity) and RS-20 (structure only) alphabets with its native spaced-seed prefilter and a candidate depth of 500. Infernal used one query-specific covariance model per RNA. BLASTN and nhmmer served as sequence-only base-lines, ranked by bit score or E-value as specified for each analysis. Software versions, commands and non-default options are provided in the Supplementary Methods.

#### Sens@1FP

It was defined as the fraction of non-self positives retrieved before the first false positive, macro-averaged across assessable queries. For Rfam annotation-defined retrieval, same-family targets were positive at the family level, while different-family targets within the same clan were neutral. At the clan level, different-family targets belonging to the same clan were positive, while same-family targets were neutral. Rfam-unassigned targets were neutral in both evaluations, and all other annotated non-positive targets were treated as false positives. Queries without a non-self positive were excluded.

#### Structure-defined ground truths

Structural relatedness was evaluated using two complementary similarity measures. Under US-align-defined truth, query–target pairs with symmetric TM-score *τ* , where *τ* 0.40, 0.45, 0.50 , were labelled positive. Under RMalign-defined truth, RMscore thresholds were selected to match the number of positive query–target pairs at each TM-score threshold. For all-neighbour retrieval, structurally similar targets were positive regardless of Rfam family. For beyond-family retrieval, same-family targets were neutral, and positives were restricted to structurally similar targets outside the query’s Rfam family. Pairs without a valid structural similarity score and queries without assessable positives were excluded from scoring. Exhaustive US-align and RMalign rankings were included as reference methods under their respective structure-defined ground truths.

#### Statistical analysis

Uncertainty was estimated using 2,000 hierarchical block-bootstrap replicates over Rfam families or clans, retaining all queries within sampled blocks. Blocking was used because queries from the same Rfam family or clan are not statistically independent. Resampling individual queries would therefore underestimate uncertainty. Confidence intervals were calculated from bootstrap percentiles, and paired method differences were evaluated using the same resamples. A difference was considered significant when its 95% paired-bootstrap confidence interval excluded zero.

### Retrieval performance across sequence identity levels

The effect of sequence identity on retrieval performance was evaluated on the held-out RNAsolo 2.0 test set using Sens@1FP, as described above. Pairwise sequence identity was calculated from global alignments using Biopython PairwiseAligner, as the number of identical aligned positions divided by the shorter sequence length. To establish an RNA-specific background distribution, we sampled 5,000 unrelated RNA pairs from different Rfam families and clans, restricted to lengths of 40–200 nt and length ratios of 0.8–1.25. Based on the median (60.5%) and 99th percentile (70.2%) of this distribution, same-family query–target pairs were divided into three identity ranges: 70.2–80%, 60.5–70.2% and *<* 60.5%.

Within each range, only same-family targets meeting the corresponding identity criterion were labelled positive. Other same-family targets, including those with identity 80%, together with different-family targets from the same clan and Rfam-unassigned targets, were treated as neutral. All remaining Rfam-annotated targets were negatives. The upper identity limit focused the analysis on sequence-divergent relationships; five queries from four families had no positives below 80% identity and were therefore excluded from the stratified evaluation.

BLASTN and nhmmer were ranked by increasing E-value, with unreported targets treated as unretrieved. RFoldSearch and RFoldSearch-TM results were averaged over ten seeded tie-orderings. Negative controls ranked targets by absolute length difference or cosine similarity of 3Dr-token composition vectors, or used RFoldSearch searches with randomly permuted query tokens and associated coordinates.

To evaluate retrieval beyond sequence-detectable positives, same-family targets reported by BLASTN or nhmmer at *E* 10 were reclassified as neutral without removing them from the search database. This deliberately permissive exclusion threshold also removed weak sequence-search matches and was not intended as a criterion for statistically significant homology.

For clan-level evaluation, positives were different-family targets from the same Rfam clan, while same-family targets were neutral. The sequence-based filter was applied to these clan-level positives using the same BLASTN and nhmmer cutoff. All 81 queries from eight clans retained at least one positive after filtering.

Sens@1FP was averaged over assessable queries. Uncertainty was estimated from 2,000 hierarchical block-bootstrap replicates, resampling Rfam families with replacement while retaining their constituent queries. All methods were evaluated using matched bootstrap resamples.

### Evaluation of runtime scalability

Runtime benchmarks followed the accuracy-evaluation configurations unless stated otherwise. For each corpus, a total of 300 queries were sampled with a fixed seed, with 15 queries drawn from each of 20 length strata defined at 5-percentile intervals. Within each corpus, the same queries were used for all methods and database sizes and were excluded from every nested target subset. Query sets for RhoFold+ and NuFold were sampled separately. Mean and median per-query search times were reported, with database construction and indexing measured separately.

Unless otherwise stated, all runtimes were measured on the same system (Intel Xeon Platinum 8470Q; NVIDIA RTX 4090 for GTalign). CPU methods were run with one CPU core per query (OMP NUM THREADS=1); US-align, RTM-align, RMalign and Infernal were run as up to 18 concurrent single-threaded jobs on separate cores. The three CPU coordinate aligners exhaustively searched every RhoFold+ subset, without size extrapolation. Rare RTM-align failures (at most 19 targets per query) were skipped, and these aligners were not run on NuFold.

Configurable methods retained up to 500 candidates. RFoldSearch used --top n 500 for this benchmark, and RFoldSearch-TM reranked the top 500. GTalign was also run with --speed=13 (GTalign (GPU)-fast) on both databases. Infernal was evaluated only on RhoFold+ in its accuracy-benchmark configuration and with cmsearch--rfam (Infernal-fast), using query-model-derived secondary structures. Reported times include only cmsearch, excluding per-query cmbuild and cmcalibrate -L 0.01. Infernal’s less stringent filtering below 2 Mb causes the step between the 10,000-and 20,000-structure subsets. RiboSeek used pre-encoded structures with RS-80 and RS-20 on both databases. RFoldSearch was benchmarked as released (version 1.0.2). Peak memory was recorded as the peak resident set size (VmHWM) of a single search process that loaded the full database, built its index and ran all 300 queries. Because RiboSeek (RS-20) required more than 90 GB, its full-database runs were performed on a machine with the same CPU type and 180 GB of memory.

### Evaluation on the NuFold benchmark

We sampled up to 50 Rfam-annotated NuFold [27] structures per family and added 15,320 unannotated structures. These were matched to the annotated set by organism, length bin and mean predicted Local Distance Difference Test (pLDDT) bin, allowing cross-organism sampling when necessary. The final benchmark contained 33,043 structures from 47 organisms and 2,264 Rfam families. Queries comprised 69 held-out experimental structures and 150 predicted structures. Predicted queries were sampled one per family from families with at least ten entries spanning at least two organisms.

For annotation-defined evaluation, targets without an Rfam annotation were treated as neutral. For predicted queries, all database entries sharing the query’s RNAcentral URS identifier, including the query itself, were neutral; positives shared the query’s Rfam family but came from a different organism, whereas same-family targets from the query organism were also neutral. Experimental queries were not present in the database; for these queries, targets with an exactly identical nucleotide sequence were neutral and all other same-family targets were positive. All remaining Rfam-annotated targets were negative.

For structure-defined evaluation, query-equivalent and unscored targets were treated as neutral. All other targets were classified as positive or negative according to thresholds on the symmetric US-align TM-score. For each TM-score threshold, the corresponding RMalign RMscore threshold was selected to produce the same overall positive fraction among query–target pairs scored by both methods.

RFoldSearch, RFoldSearch-TM and RiboSeek used a candidate depth of 500, and RFoldSearch-TM reranked RFoldSearch candidates by US-align TM-score. BLASTN and nhmmer hits with *E* 10 were retained. These hits and Infernal hits were ranked by bit score, and exhaustive coordinate-based aligners were ranked by symmetric structure score. Neutral targets were removed from each ranking before evaluation. Sens@1FP was calculated as the number of positives retrieved within the top 200 hits before the first false positive, divided by min(*N*_positive_, 200). Confidence intervals for means and paired differences were obtained from 2,000 Rfam-family-block bootstrap replicates.

Predicted queries were stratified by query mean pLDDT (*<* 70, 70–79, 80–89 and 90), and experimental queries by target mean pLDDT. In sequence-filtered analyses, positives detected by BLASTN or nhmmer at *E* 10 were reclassified as neutral. For the top-200 recovery analysis, sequence-undetected family members were same-family targets missed by BLASTN and nhmmer at *E <* 0.01 and absent from the Infernal ranking. Sampling strata and filtering thresholds were prespecified.

### Reference-based evaluation of nucleotide-level alignment accuracy

Nucleotide-level alignment accuracy was evaluated against Rfam 3D seed alignments mapped to experimentally determined PDB structures. PDB entries used for RFold-Search training or validation were excluded. For each seed-alignment row, we retained at most one structure with at least 70% sequence coverage and sequence identity to the resolved chain. Within-family pairs were constructed from independently delimited Rfam regions with insertions retained. Each pair was required to share at least 20 assessable consensus match-state columns. The final benchmark comprised 313 pairs from 125 structures across 24 families.

Reference correspondences were defined as nucleotide pairs occupying the same Rfam consensus match-state column. Only columns with valid mapped positions for all compared structural methods were evaluated. Insert states, gaps and other non-assessable positions were excluded. Rfam was used only to delimit regions and define reference correspondences after alignment, and was not provided as input to any method.

RFoldSearch aligned 3Dr tokens by Smith–Waterman local alignment with affine gap penalties and the Structure-BLOSUM matrix. RiboSeek used RS-20 with its native substitution matrix and local alignment with a linear gap score of 2.0. US-align was run in RNA mode. As a sequence baseline, the same regions were globally aligned with Biopython PairwiseAligner (version 1.85) [43], using match, mismatch, gap-opening and gap-extension scores of +2, −1, −3 and −0.5, respectively. All predicted correspondences were mapped back to the original nucleotide indices before evaluation.

Accuracy was quantified by precision, recall and *F*_1_ under two criteria: exact pair matching and a tolerant criterion allowing an offset of up to one nucleotide in each structure. Predicted pairs involving Rfam insert states were treated as neutral, and pairs without an alignment were assigned scores of zero. The primary summary statistic was family-macro *F*_1_, obtained by averaging pair-level scores within each family and then across the 24 families. Confidence intervals for each method and for paired differences between methods were derived from 2,000 family-block bootstrap replicates, using the same resamples for all methods.

### Recurrent token-word detection and structural annotation

We defined recurrent four-token 3Dr words as those occurring at least eight times across at least five chains, three Rfam families and two Rfam clans in the curated RNAsolo 2.0 set. Both RMSD analyses used the same four-token windows from Rfam-labelled chains, restricted to occurrences with unique mappings to the deposited structure and complete base-inclusive coordinates. The same sampled occurrence pairs were evaluated using the five descriptor anchors and a base-inclusive atom correspondence comprising all backbone and sugar heavy atoms plus three base atoms: N9, C4 and C8 for purines, or N1, C2 and C6 for pyrimidines. This correspondence allowed superposition across different nucleotide sequences. Fragments were superposed by Kabsch alignment and summarised by median pairwise RMSD [44, 45]. Only crossclan occurrence pairs were used, treating each family without a clan assignment as a separate clan, with at most 400 pairs sampled per word.

Candidates were compared with frequency-matched non-candidate words, sequence-matched windows and random windows. Frequency-matched controls occurred at least eight times in the shared set, contained at least one cross-clan pair and were binned by occurrence count (8–15, 16–31 or 32); candidates with fewer than eight occurrences were assigned to the 8–15 bin. For display, each candidate was compared with the median of 25 controls sampled from its bin. Sequence-matched windows shared the nucleotide sequence of each candidate occurrence but not its token word, and random windows were unconstrained; both were evaluated in 25 replicates of at most 25 cross-clan pairs. Candidates and frequency-matched controls were compared for each atom set by permuting unique-word labels 10,000 times within bins (two-sided, plus-one correction).

Words were classified as standard helical when their modal Biotite pairing pattern placed all four nucleotides in cis pairs with consecutive antiparallel partners, and as other words otherwise. BGSU hairpin and internal-loop overlap [46, 47] required at least one shared nucleotide. Each of 1,000 null replicates assigned every word the same number of random four-nucleotide windows from BGSU-covered chains. Median per-word overlap fractions were compared with the null overall and by word class (one-sided, plus-one correction).

For motif-specific examples, token positions were mapped to BGSU nucleotides by PDB entry, model, chain, nucleotide identity, author residue number and insertion code. A window was considered contained in a motif when all its nucleotides belonged to the same annotated motif occurrence. GNRA-associated fragments were six-token instances of 44-48-62-71-66-12 covering all six aligned nucleotides of an HL 34789.6 occurrence, and T-loop fragments were occurrences of 48-24-39-39 contained within HL 33597.6, covering four of its nine aligned nucleotides. Fragments were selected independently of DSSR annotations.

DSSR (version 2.9.2) [28] was used to annotate Leontis–Westhof interactions [29], retaining all pairing records, including those with partners outside the token window. Labels were ordered from each focal nucleotide towards its partner; the two edge letters were exchanged when the focal nucleotide was the second member of the DSSR pair record. A fragment matched a positional signature when every required position had at least one pairing record with the specified label; other positions were unconstrained, and additional pairings were permitted. Residue labels in the displayed T-loop representative and its external partners follow deposited numbering. Structural views use the base-inclusive superpositions, with the same transformations applied to displayed external partners; these partners were not included in the fit.

### E-values

E-values were calibrated separately for experimental (represented by RNAsolo 2.0 [4]) and predicted (represented by NuFold [27]) structure databases in target-cluster space, with targets clustered at 80% nucleotide-sequence identity using CD-HIT-EST [32] to account for database redundancy.

Let *S*(*q, j*) denote the token Smith–Waterman score used to estimate significance for query *q* and target *j*. Experimental structures used raw scores, whereas predicted structures used composition-adjusted substitution scores based on the surrounding query tokens. Each non-empty target cluster contributed its maximum eligible score,

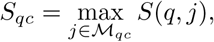

where *M*_qc_ excludes the query’s own cluster, members sharing a recorded Rfam family or clan, and members identified as structurally related by US-align. Structural auditing, member exclusion and cluster-leader replacement were performed iteratively within the calibration range. No query–target length-ratio filter was applied.

Cluster maxima were modelled by a target-aware Gumbel distribution,

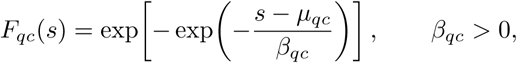

with

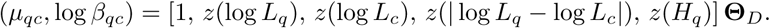

Here, *L_q_* is query-token length, *L_c_* is median cluster-member token length, *H_q_* is query-token entropy, and *z* denotes standardisation using the weighted Gumbel-training observations. Parameters were fitted by regularised weighted likelihood, with left-censoring used by default to emphasise the upper score tail.

For a candidate score *s*, the initial and calibrated E-values were

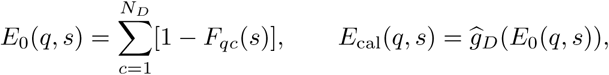

where *N_D_* is the number of target clusters and *g_D_* is a non-decreasing empirical map fitted to family-balanced background counts. Calibration and sensitivity were evaluated on held-out Rfam families and clans. Background auditing, fitting, extrapolation and validation are described in the Supplementary Methods (E-value calibration details).

### Web server and API

RFoldSearch is available at https://www.rfoldsearch.com and supports three RNA structure databases. PDB-RNA contains all PDB chains classified as RNA polymer entities by the RCSB PDB [48] (18,909 chains; built on 21 September 2026). RNA-solo 2.0 contains representative RNA chains from the BGSU non-redundant list [4] (14,531 chains; release 4.60). NuFold RNA Structure Database contains predicted RNA structures [27] (514,123 structures; version 9-14-2026). This is a later release than the one used for the runtime benchmarks (version 1-13-2026; 531,888 structures after length filtering).

Token databases and precomputed *k*-mer indices are retained in memory. Searches run on a single CPU core using RFoldSearch or RFoldSearch-TM, with 50 hits displayed per database by default. Users can upload or paste PDB/mmCIF coordinates and adjust the number of displayed hits, candidate depth and prefilter threshold. RFoldSearch ranks candidates by the structure score, whereas RFoldSearch-TM reranks candidates using US-align.

Depending on the search mode, results include structure scores, TM-scores, RMSD, alignment coverage, nucleotide or 3Dr-token identity, and calibrated E-values. E-values are reported only for the default RFoldSearch mode. Experimental structures are linked to PDB metadata, while predicted structures include RNAcentral identifiers, organism information, available Rfam annotations and model-confidence metrics. Pairwise alignments and superposed structures are displayed interactively using Mol* [49], with 3Dmol.js [50] as a fallback for browsers without WebGL2. Results can be shared by URL and downloaded as TSV or JSON.

The server is implemented using FastAPI and requires no registration. A REST API provides programmatic access to structure searches, results, alignments and superposed coordinates. Interactive API documentation is available at https://www.rfoldsearch.com/api/docs.

### Use of AI tools

Analysis and plotting code was developed with the assistance of AI coding assistants (OpenAI Codex and Anthropic Claude). All code was reviewed, run and validated by the authors, who take full responsibility for the analyses and results reported here.

## Data availability

The processed datasets generated and analysed in this study, including data underlying the benchmarking results and figures, are available at https://github.com/Cassie818/ RFoldSearch-analysis. The source RNA structures and annotations were obtained from the publicly available databases described in the Methods.

## Code availability

RFoldSearch is freely available as open-source software. The source code is available at https://github.com/DylanYang/RFoldSearch. The version described here (version 1.0.2) is archived at Zenodo (https://doi.org/10.5281/zenodo.24412569). The RFold-Search web server is accessible at https://www.rfoldsearch.com/, and its source code is available in the same repository. Scripts used to reproduce the benchmarking analyses and figures presented in this study are available at https://github.com/Cassie818/ RFoldSearch-analysis.

## Acknowledgements

Figure 1 was prepared using BioRender.com. R.C. was supported by an Australian Government Research Training Program Scholarship and the M.G. & R.A. Plowman Scholarship in Inorganic Chemistry. H.L. was supported by the Yunnan Key Laboratory of Modern Analytical Mathematics and Applications (No. 202302AN360007) and the Yunnan Cross-integration Innovation Team of Modern Applied Mathematics and Life Sciences (No. 202405AS350003).

## Author contributions

R.C., M.B. and M.H. conceived the study. R.C. designed and developed the structural alphabet and the RFoldSearch search framework, and carried out the main computational experiments. W.Y. implemented components of the software, conducted runtime benchmarks and developed the web server. H.L. worked on statistical modelling and quantitative evaluation. B.R., M.B. and M.H. provided scientific guidance and contributed to the interpretation of the findings. R.C. prepared the initial manuscript, which was reviewed and edited by all authors. M.B. and M.H. supervised the project.

