## Supplementary material for "RFoldSearch enables fast RNA structure search beyond sequence similarity": supp files

### Supplementary Materials

#### Supplementary Methods

##### Descriptors

The final 17-dimensional descriptor comprises the first spatial-neighbour FRAME block $\mathbf{g}_{\text{nn}}^{(1)}$ and the context vector $\mathbf{g}_{\text{ctx}}$.

*Virtual centre for neighbour selection.* Spatial neighbours are selected using an RNA virtual centre inspired by Foldseek [18], favouring base-proximal contacts over sugar–backbone proximity alone. Let $\mathbf{x}_i^B$ be the glycosidic nitrogen (N9 for purines and N1 for pyrimidines) and $\mathbf{o}_i$ the frame origin. We define

$$\mathbf{u}_i = \frac{\mathbf{x}_i^B - \mathbf{o}_i}{\|\mathbf{x}_i^B - \mathbf{o}_i\|}, \qquad \mathbf{v}_i = \frac{\mathbf{o}_{i+1} - \mathbf{o}_i}{\|\mathbf{o}_{i+1} - \mathbf{o}_i\|}, \qquad \mathbf{w}_i = \frac{\mathbf{u}_i \times \mathbf{v}_i}{\|\mathbf{u}_i \times \mathbf{v}_i\|},$$

with virtual centre

$$\mathbf{c}_i = \mathbf{o}_i + \alpha \mathbf{u}_i + \beta \mathbf{v}_i + \gamma \mathbf{w}_i.$$

We used the untuned symmetric setting $\alpha = \beta = \gamma = 0.5$.

For feature-ablation experiments (Fig. S2), we additionally evaluated three FRAME-based blocks: a second nonlocal spatial neighbour $\mathbf{g}_{\text{nn}}^{(2)} \in \mathbb{R}^{10}$, a base-pair partner $(\mathbf{g}_{\text{bp}}, m_{\text{bp}}) \in \mathbb{R}^{11}$ and a stacking partner $(\mathbf{g}_{\text{st}}, m_{\text{st}}) \in \mathbb{R}^{11}$. The latter two apply the same FRAME template to the annotated interaction partner and include a binary availability mask and FRAME values were set to zero when no partner was available. Their normalisation statistics were estimated only from nucleotides with active masks, whereas mask dimensions were left unnormalised. All other dimensions were z-scored using training-set statistics.

The full ablation descriptor was therefore

$$\mathbf{x}_i^{\text{full}} = \left[\mathbf{g}_{\text{nn}}^{(1)}, \mathbf{g}_{\text{nn}}^{(2)}, \mathbf{g}_{\text{bp}}, m_{\text{bp}}, \mathbf{g}_{\text{st}}, m_{\text{st}}, \mathbf{g}_{\text{ctx}}\right] \in \mathbb{R}^{49}.$$

The additional 32 dimensions were used only for feature ablation and were not included in the final RFoldSearch descriptor.

##### Substitution matrix

The deployed RFoldSearch model uses Structure-BLOSUM (see "Methods – 3Dr substitution matrix"), derived from token co-occurrences between structurally similar training chains. We compared it with four alternatives under identical retrieval settings (Fig. S5).

*Identity matrix.* Exact token matches scored +2 and mismatches −1.

*Codebook-cosine matrix.* Token pairs were scored by the cosine similarity of their L2-normalised codebook embeddings, $S(i,j) = \mathbf{e}_i^\top \mathbf{e}_j$.

*Codebook-BLOSUM matrix.* Codebook vectors were normalised to unit length, and their cosine similarities were converted to joint probabilities by $p_{ij} \propto \exp(\hat{\mathbf{e}}_i^\top \hat{\mathbf{e}}_j / \tau)$ with $\tau = 0.5$. The matrix was scored as $2 \log_2[p_{ij}/(p_i p_j)]$ with marginals $p_i = \sum_j p_{ij}$ of this joint distribution, and the diagonal was rescaled to unit mean.

*Rfam-BLOSUM matrix.* For each Rfam family with at least two training chains, at most 100 unordered chain pairs were sampled. Their token strings were aligned by Smith–Waterman with an identity scoring matrix and affine gap penalties of $-11$ for gap opening and $-1$ for gap extension. Every non-gap aligned column added one count to both $C_{ij}$ and $C_{ji}$, and the resulting log-odds matrix $2\log_2[p_{ij}/(p_i p_j)]$ was rescaled to unit mean diagonal.

#### Evaluation metrics

In addition to Sens@1FP, ranked retrieval quality was evaluated using Recall@1 and mean average precision.

*Recall@1.* Recall@1 was the fraction of queries whose top-ranked assessable non-self target was a true positive, with neutral targets skipped.

*MAP.* Average precision (AP) was computed from each query's ranked hit list and averaged equally across queries to obtain MAP. Self-hits and neutral targets were removed before assigning effective ranks, whereas unreturned positives remained false negatives. Macro-averaged precision–recall curves were computed by averaging interpolated precision across queries on a common recall grid. Each method used its native within-query ranking.

*AUROC.* For each query, AUROC was calculated as the probability that a randomly selected positive was ranked ahead of a randomly selected negative, after excluding the query itself and neutral targets. Targets not returned by a method were treated as tied below all returned targets; consequently, an unreturned positive–negative pair contributed 0.5. Per-query AUROC values were macro-averaged across queries.

#### Structure-score refinement details

Candidates passing the $k$-mer prefilter were aligned to the query by affine-gap token Smith–Waterman, with the traceback obtained in the same dynamic-programming pass as the alignment score $S_{\mathrm{SW}}$. The resulting nucleotide correspondences were evaluated in Cartesian space using stored C4′ coordinates, without re-optimising the alignment.

For a Smith–Waterman traceback containing $L_{\mathrm{aln}}$ non-gap nucleotide pairs, let $L_Q$ and $L_T$ denote the numbers of tokenised nucleotides with valid coordinates in the query and target, respectively, and let $r_i$ denote the C4′–C4′ distance of aligned pair $i$ after superposition. We defined

$$T_X = \frac{1}{L_X}\sum_{i=1}^{L_{\mathrm{aln}}} \frac{1}{1+[r_i/d_0(L_X)]^2}, \qquad X \in \{Q, T\},$$

with

$$d_0(L) = \max\left\{0.5,\ 0.6\sqrt{L-0.5}-2.5\right\}\ \text{Å},$$

and the alignment TM-score

$$\mathrm{TM}_a = \frac{1}{2}(T_Q + T_T).$$

Normalisation by both chain lengths penalises partial coverage.

The superposition was optimised by iterative Kabsch refinement from two initialisations: all aligned pairs and the highest-scoring gapless segment under Structure-BLOSUM. At each iteration, a Kabsch transformation fitted to the current inliers was applied to all aligned pairs to calculate $\mathrm{TM}_a$. Inliers were then updated to pairs satisfying

$$r_i < \min\{d_0(L_Q), d_0(L_T)\} + 1\,\text{Å}.$$

Refinement continued for at most five iterations and stopped earlier if the inlier set was unchanged or fewer than three inliers remained. The highest $\mathrm{TM}_a$ obtained across both initialisations and all iterations was retained. Alignments containing fewer than three nucleotide pairs were assigned $\mathrm{TM}_a = 0$.

Hits were ranked by the structure score

$$R_{\mathrm{str}} = S_{\mathrm{SW}} \times \mathrm{TM}_a,$$

which combines token-level similarity with the global spatial consistency of the aligned nucleotides. Ranking ties were resolved using the same procedure as at the prefilter boundary.

#### E-value calibration details

*Scores and background construction.* Experimental structures used raw token Smith–Waterman scores. For predicted structures, alignment used composition-adjusted substitution scores,

$$P_{ia} = M(q_i, a) - \left[\frac{1}{|W_i|}\sum_{k\in W_i} M(q_k, a) - \sum_b \pi_D(b)M(b, a)\right],$$

where $M$ is the Structure-BLOSUM substitution matrix, $W_i$ contains positions within 20 tokens of query position $i$ excluding $i$, and $\pi_D$ is the target-database token composition. The resulting alignment score was denoted by $S(q, j)$. Composition adjustment affected E-value estimation only; structure-score ranking retained the unadjusted score $S_{\mathrm{SW}}$.

The query's own cluster was excluded. Within the remaining clusters, members were excluded if they shared an Rfam family or clan with the query according to the annotations used for calibration, or if their mean query- and target-normalised US-align TM-score was at least 0.45. Cluster maxima were recomputed after exclusion, allowing another eligible member to replace an excluded leader. No query-target length-ratio filter was applied.

Structural auditing used a database-specific initial E-value ceiling $\tau_D$. Unaudited cluster leaders with $E_0 \leq \tau_D$ were aligned using US-align, together with all other unaudited eligible members of the same cluster satisfying this threshold under the current model. Related members were excluded, cluster leaders were recomputed, and the Gumbel model was refitted until no qualifying unaudited leaders remained. The same initial E-value ceiling, $\tau_D$, was used for fitting the empirical map and was recorded with the fitted parameters. Background pairs outside this range were not exhaustively screened.

*Gumbel fitting.* Query-token Shannon entropy $H_q$ was calculated using natural logarithms. Features were standardised using the weighted mean and standard deviation of the Gumbel-training observations. Parameters were fitted by weighted likelihood with an $L_2$ penalty on non-intercept coefficients.

By default, scores at or below the weighted 95th percentile were left-censored within strata defined by $\min(L_q, L_c)$ and the query–cluster length ratio. Higher scores contributed their observed values, whereas censored observations contributed the Gumbel cumulative probability at the threshold. Strata containing fewer than 200 observations used the global weighted threshold. Experimental fitting also supported an uncensored likelihood.

Cluster maxima shared each query's weight equally. Experimental queries were weighted equally within Rfam families and families equally. Predicted fitting assigned 80% of the training weight to core queries and 20% to auxiliary queries, with equal query weights within each panel. A separate annotated query panel was used for empirical calibration.

*Empirical calibration and extrapolation.* Experimental calibration maxima were recomputed over eligible annotated members; predicted calibration used all eligible members. Denoting these maxima by $S_{qc}^{\text{cal}}$, we calculated

$$E_{0,qc} = E_0(q, S_{qc}^{\text{cal}}).$$

At each observed $E_0$ knot $e_k$, the empirical map was defined by the family-balanced cumulative background count,

$$\widehat{g}_D(e_k) = \sum_{f\in\mathcal{F}_D} \sum_{q\in Q_f} \sum_{c\in\mathcal{B}_q} \frac{\rho_q}{|\mathcal{F}_D|\,|Q_f|} \mathbf{1}(E_{0,qc} \le e_k),$$

where $\mathcal{F}_D$ contains the calibration-query families, $Q_f$ contains queries assigned to family $f$, and $\mathcal{B}_q$ contains retained background clusters with $0 < E_{0,qc} \le \tau_D$.

Experimental maps used the annotated fitting queries, with $\rho_q = N_D/B_q$ scaling the annotated background to the full target-cluster space. Here, $B_q$ was counted after own-cluster and family/clan exclusions but before structural exclusion. Predicted maps used the separate annotated calibration panel with $\rho_q = 1$, because all eligible target clusters were evaluated.

The map was linearly interpolated between knots. A positive power-law exponent was fitted by least-squares regression of log cumulative background counts on log $E_0$, using logarithmically spaced thresholds with positive counts within the fitting range. Extrapolation below and above the observed range used this exponent, anchored at the corresponding endpoint, with E-values capped at $N_D$. Extrapolated values were not considered independently validated estimates.

*Full-database NuFold calibration.* In addition to the 33,043-structure analysis shown in Fig. S15, calibration was performed on the NuFold database served by the web server (version 9-14-2026), containing 514,123 chains in 181,082 sequence clusters. This analysis used 600 core queries, 150 auxiliary short or low-entropy queries, and 600 annotated calibration queries. The panels were chain- and cluster-disjoint, and the core and calibration panels shared no Rfam family or clan. Core queries represented distinct families and were balanced across length deciles. Structural auditing and empirical fitting used $\tau_D = 400$.

*Validation.* Calibration and sensitivity were evaluated on held-out Rfam families and clans. RNAsolo used fully audited annotated backgrounds, whereas PDB-RNA used existing US-align adjudications of high-scoring pairs. For the full-database NuFold analysis, the Gumbel model fitted on the separate training panel was held fixed. Each calibration query was evaluated using an empirical map refitted after excluding that query and all calibration queries sharing its family or clan.

Calibration compared family-balanced background-cluster counts with reported E-value thresholds. Sensitivity measured the fraction of same-family target clusters retrieved. NuFold confidence intervals were estimated by bootstrap resampling of calibration queries.

#### Retrieval performance under alternative data splits

We performed five-fold cross-validation within the original training–validation pool, excluding the independent test partition. All schemes used 237 labelled queries of at most 500 nucleotides from 41 Rfam families with at least two chains each. Family-disjoint and homology-disjoint schemes shared family-grouped folds, approximately balanced by query count. RF00005 contained 104 queries and formed one fold. PDB-disjoint folds grouped chains by PDB entry.

Relatedness to held-out queries was defined by symmetric US-align TM-score $\geq 0.45$ or BLASTN/nhmmer $E < 0.01$ in either direction. Unlabelled chains assigned to held-out families by `cmscan` at $E < 0.1$ were also considered related. Family-disjoint filtering removed held-out-family chains and related unlabelled chains. Homology-disjoint filtering additionally removed related labelled chains and labelled members of the held-out clans. A matched PDB-disjoint control randomly removed the same fractions of labelled and unlabelled training chains as the corresponding homology-disjoint fold.

The remaining chains were split 80:20 for training and validation. Descriptor normalisation, VQ-VAE fitting and Structure-BLOSUM estimation used only the training subset. Structure-BLOSUM used within-training US-align pairs with symmetric TM-score $\geq 0.5$. Validation guided model selection, and all hyperparameters matched the final RFoldSearch model.

Held-out queries were searched against the same complete structure database across all partitioning schemes, including structures excluded from model training.

This kept the retrieval targets and relevance labels fixed, allowing us to isolate the effect of training-set relatedness on retrieval performance. Self-hits were excluded, but filtered training chains remained eligible targets, keeping the database and relevance labels fixed. Each scheme used five independent VQ-VAE initialisations. Performance was averaged over three seeded hash orderings that resolved ties at the prefilter cutoff and in the final ranking.

#### Runtime summary statistics

Search time was measured as wall-clock time for individual queries against prepared databases. CPU methods used one CPU core per query, whereas GTalign used one GPU. One-off database preparation was excluded. For each method and database size, we calculated the mean and median search times over the same 300 queries, together with the 25th and 75th percentiles for RFoldSearch.

In Figs. 3A and B, mean per-query search times were plotted directly against database size without per-target normalisation. For full-database comparisons, mean query times were also expressed relative to RFoldSearch on the same database and query set. To compare scaling independently of absolute runtime, the median (Figs. S13A and B) or mean (Figs. S13C and D) query time was divided by $N/1{,}000$, where $N$ is the number of target structures, and then normalised to its value at $N = 1{,}000$. RFoldSearch's interquartile bounds were normalised separately. Scaling exponents were estimated by ordinary least-squares regression of $\log t$ on $\log N$ across eight NuFold database sizes, where $t$ is the mean query time underlying Fig. S17A. RiboSeek/RFoldSearch runtime ratios were calculated from the mean query times at each database size.

To characterise query-level variability, per-query speed-ups were calculated as $t_{\text{RiboSeek}}/t_{\text{RFoldSearch}}$ for RiboSeek (RS-20) on the full NuFold database (Fig. S17B). We reported the number of queries with a speed-up above one and the median speed-up across the 300 queries. Query-length dependence was assessed by ordinary least-squares regression of $\log t$ on $\log L$, where $L$ is the number of 3Dr tokens, with goodness of fit reported as $r^2$ (Figs. S13E and F). Trend lines connect median times in eight logarithmically spaced length bins, excluding bins containing fewer than five queries. The contribution of the slowest queries was calculated as the fraction of total runtime accounted for by the 30 slowest of the 300 queries.

#### Benchmark tool versions and command lines

RFoldSearch and its TM-reranking mode, RFoldSearch-TM, were benchmarked against four coordinate-based aligners (RMalign [13], US-align [12], RTM-align [51] and GTalign [52]), the structural-alphabet method RiboSeek [22], the sequence-and-secondary-structure profile method Infernal [53], and the sequence-based methods BLASTN [15] and nhmmer [16]. For accuracy evaluation, each query was searched against the full RNAsolo database, with targets ranked by each method's native pairwise or combined score.

#### RFoldSearch

RFoldSearch (version 1.0.2) encoded each chain as a sequence over the 80-state 3Dr alphabet and ran the deployed pipeline: an exact 4-mer inverted-index prefilter with one-substitution Structure-BLOSUM expansion, followed by affine-gap token Smith–Waterman alignment of the top 200 candidates and ranking by structure score (Table S9).

```
rfoldsearch search --query chain_id --db target_db.pt --top_k 10
```

#### RFoldSearch-TM

RFoldSearch-TM (version 1.0.2) is an accuracy-oriented mode that keeps the same prefilter and candidate pool but realigns retained candidates with US-align and ranks them by symmetric TM-score. It gives up the speed of token alignment and pair refinement for direct structural comparison, reports TM-scores but no E-values, and requires a US-align executable. Its results are reported alongside RFoldSearch wherever the two differ.

```
rfoldsearch search --query chain_id --db target_db.pt --top_k 10 --rerank tm
```

#### RMalign

RMalign [13] (version 1.1) was run as an RNA-specific sequential structural aligner. Because RNAalign requires legacy PDB input, mmCIF chains were converted to PDB format with Biopython [43]. For each pair, targets were ranked by the symmetric RMscore, defined as the mean of the scores normalised by the query and target lengths.

```
RNAalign -A query.pdb -B target.pdb
```

#### US-align

US-align [12] (version 20220924) was run in RNA mode with tabular output. Targets were ranked by the symmetric TM-score, defined as the mean of the query- and target-normalised scores.

```
USalign query.cif target.cif -mol RNA -outfmt 2
```

#### RTM-align

RTM-align [51] was run in RNA mode and ranked by the symmetric RTM-score, matching the convention used for the other coordinate aligners.

```
RTMalign query -dir2 targets/ list -suffix .cif \
         -mol RNA -outfmt 2 > out.tab
```

#### GTalign

GTalign [14, 52] (version 1.0.1) was run with disabled prefilters and ranked by symmetric TM-score. Its length filters skipped 3 of the 885 database chains, while all family-assessable and clan-assessable queries were retained.

```
gtalign --qrs=queries --rfs=targets \
        --pre-score=0 --pre-similarity=0 --nhits=885 \
        --speed=8 -o out
```

##### RiboSeek

RiboSeek [22] (version 0.2.1) was run with both of its alphabets on a database encoded with its own feature extractor under default settings. Searches used its default spaced-seed prefilter, which passes the 500 highest-scoring chains (`prefilter_candidates`=500) to global and local alignment rescoring. Because RiboSeek ranks all retained candidates internally and `top_n` controls only how many are returned, we set `top_n`=500 to recover the full ranking. When fewer than 500 chains share a seed with the query, the remaining candidate slots are filled with chains of zero prefilter score in database order; these are aligned and ranked like any other candidate.

```
Searcher(Alphabet.from_pretrained("rs80"|"sa20"),
         db, gap_penalty=-2.0).search(
         query, top_n=500,
         prefilter=True,
         prefilter_candidates=500)
```

##### Infernal

Infernal [53] (version 1.1.5) was run with one query-specific covariance model per RNA. Query secondary structure was derived from the 3D coordinates using Watson–Crick and wobble base pairs, reduced to a nested structure and encoded in a single-sequence Stockholm alignment with `SS_cons`. Models were built with `cmbuild`, calibrated with `cmcalibrate -L 0.01` and searched with `cmsearch`.

```
cmbuild -F query.cm query.sto
cmcalibrate -L 0.01 query.cm
cmsearch --toponly -T 0 --tblout out.tbl query.cm db.fasta
```

##### BLASTN

BLASTN from BLAST+ [15] (version 2.17.0) served as the sequence-only baseline. A nucleotide database was built from all chain sequences, and each query was searched with `-max_target_seqs` set to the database size. Self-hits were excluded, and targets were ranked by descending bit score.

```
blastn -query q.fasta -db db -evalue 10
       -outfmt 6 -max_target_seqs 885
```

##### nhmmer

nhmmer from HMMER [16] (version 3.4) performed profile-based nucleotide homology search. Each query sequence was searched in single-query mode against the same nucleotide database, restricted to the sense strand and the RNA alphabet. The best bit score was kept per target, self-hits were excluded and targets were ranked by descending bit score.

```
nhmmer --watson --rna -E 10 --noali
        --tblout out.tbl  q.fasta  db
```

#### Supplementary Figures

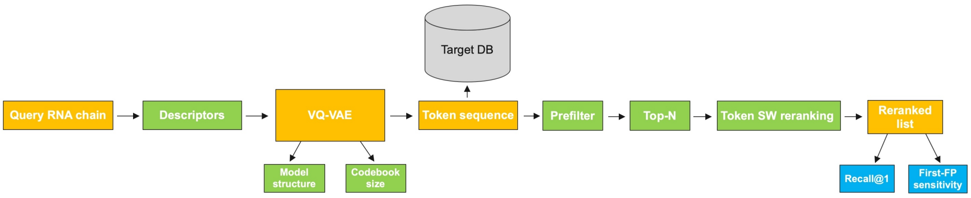


**Fig. S1**: **Evaluation framework for ablation experiments.**
Each RNA structure is encoded as a discrete token sequence and searched against a tokenised target database. Feature-group ablations (Fig. S2) use $k$-means to isolate descriptor effects from tokeniser stochasticity, whereas VQ-VAE architecture and codebook-size ablations (Figs. S3 and S4) use VQ-VAE. Green modules indicate components varied across experiments, including descriptor composition, tokenisation, prefiltering and token alignment. Retrieval was evaluated by family- and clan-level Sens@1FP over five PDB-disjoint cross-validation folds of the train–validation partition (mean ± s.d.). The selected settings were used to construct the final RFoldSearch model.

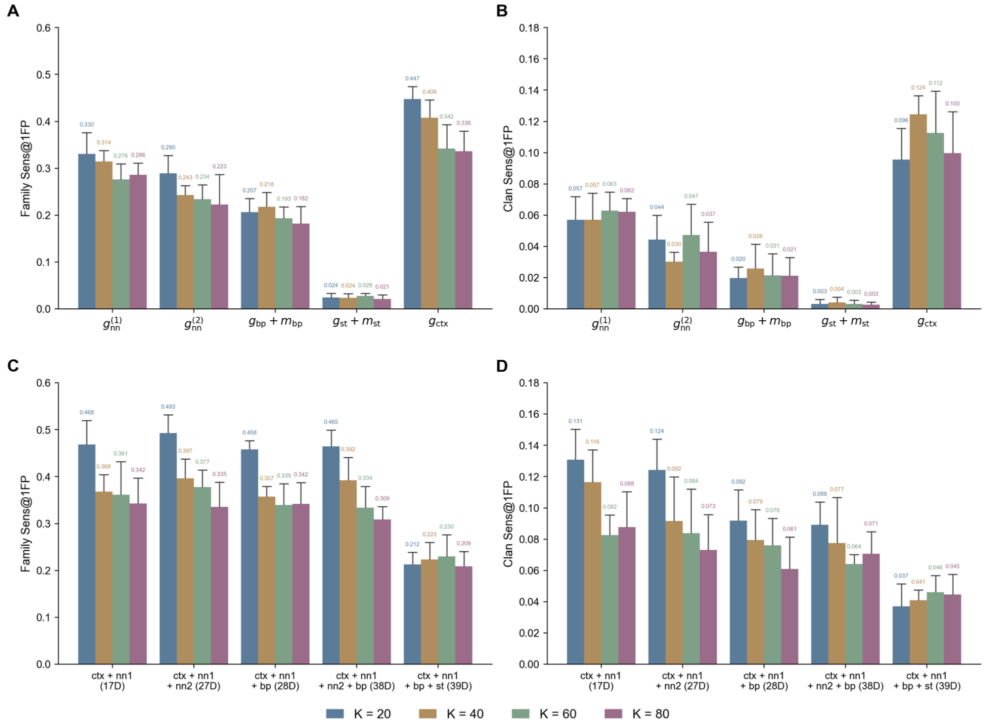


**Fig. S2**: **Feature-group ablation across codebook sizes.**
(**A**, **C**) Family-level and (**B**, **D**) clan-level Sens@1FP for (**A**, **B**) single feature groups and (**C**, **D**) feature-group combinations, evaluated by PDB-disjoint 5-fold cross-validation on the train–validation partition with the fixed retrieval settings in Table S2. Bars show means and error bars standard deviations across folds.

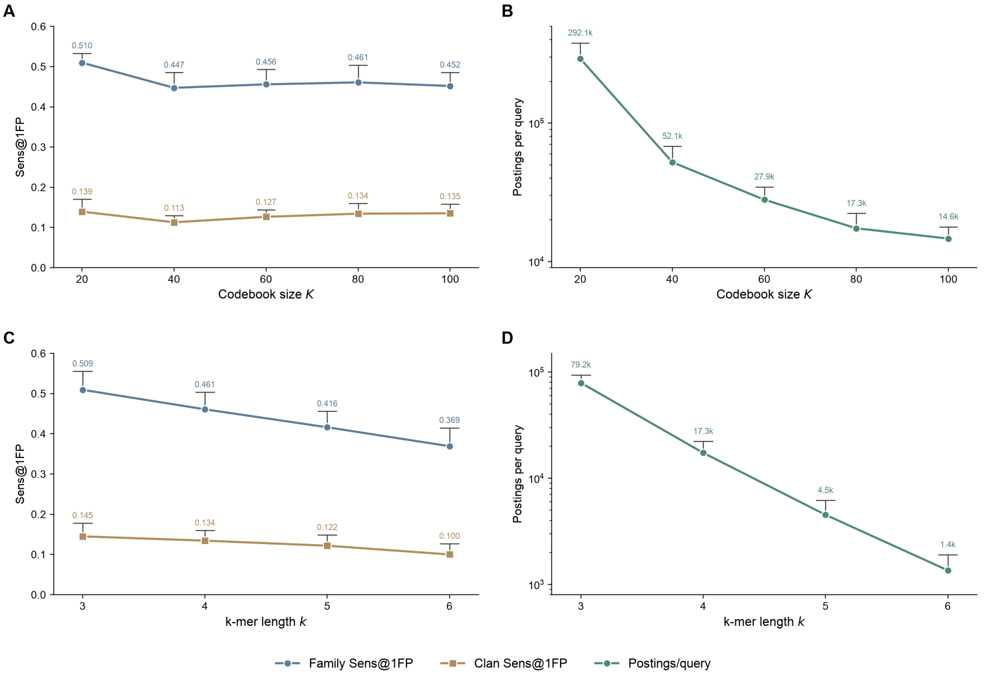


**Fig. S3**: **Codebook size and $k$-mer length trade retrieval sensitivity for prefilter selectivity.**
**(A, B)** Family- and clan-level Sens@1FP (A) and mean postings traversed per query (B) for VQ-VAE models trained on the 17-D descriptor with $K = 20$–$100$ at fixed $k = 4$. **(C, D)** The same metrics for $k = 3$–$6$ at fixed $K = 80$. Postings (log scale) are counted after similar-$k$-mer expansion but before the double-diagonal filter and candidate cutoff. All experiments use codebook-cosine substitution scores and the retrieval settings in Table S3. Mean ± s.d. over five PDB-disjoint folds.

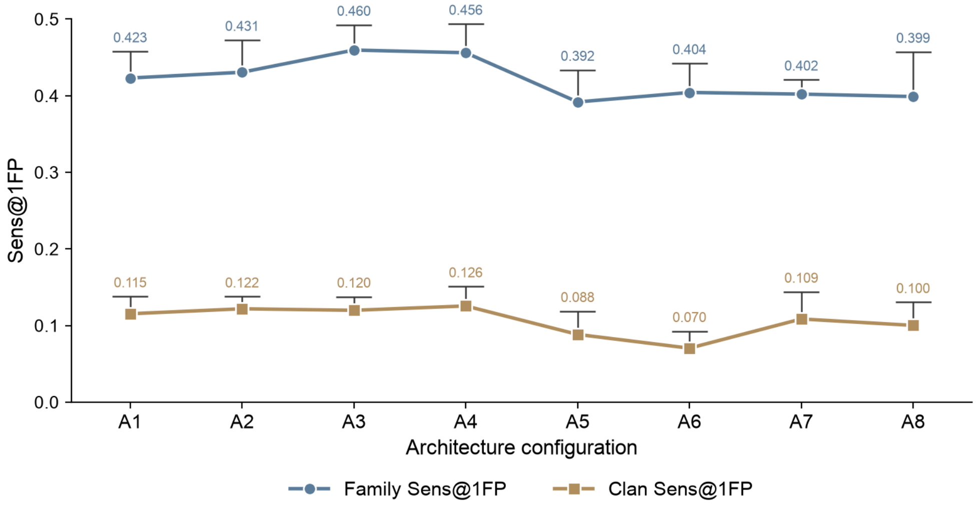


**Fig. S4**: **VQ-VAE architecture ablation on the 17-D descriptor.** Family-level and clan-level Sens@1FP for the eight different configurations from Table S5, evaluated by PDB-disjoint 5-fold cross-validation on the train–validation partition. All experiments use the fixed retrieval settings specified in Table S4. Points show the mean and error bars the standard deviation across the five folds.

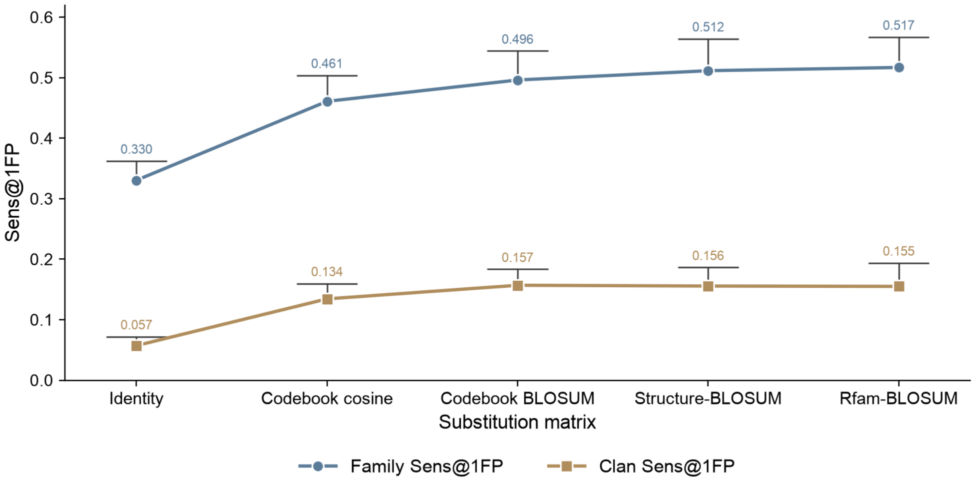


**Fig. S5**: **Substitution-matrix ablation for token Smith–Waterman reranking.**
Family-level and clan-level Sens@1FP for five candidate matrices, evaluated by PDB-disjoint 5-fold cross-validation on the train–validation partition. All experiments use the fixed retrieval settings specified in Table S6. Points show the mean and error bars the standard deviation across the five folds.

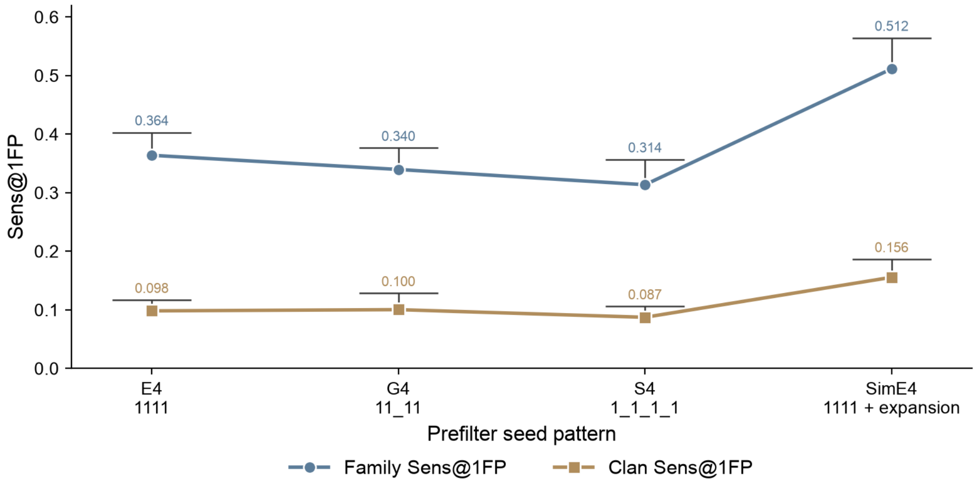


**Fig. S6**: **Prefilter seed-pattern ablation.**
Family-level and clan-level Sens@1FP for four candidate seed patterns, evaluated by PDB-disjoint 5-fold cross-validation on the train–validation partition. All experiments use the fixed retrieval settings specified in Table S7. Points show the mean and error bars the standard deviation across the five folds.

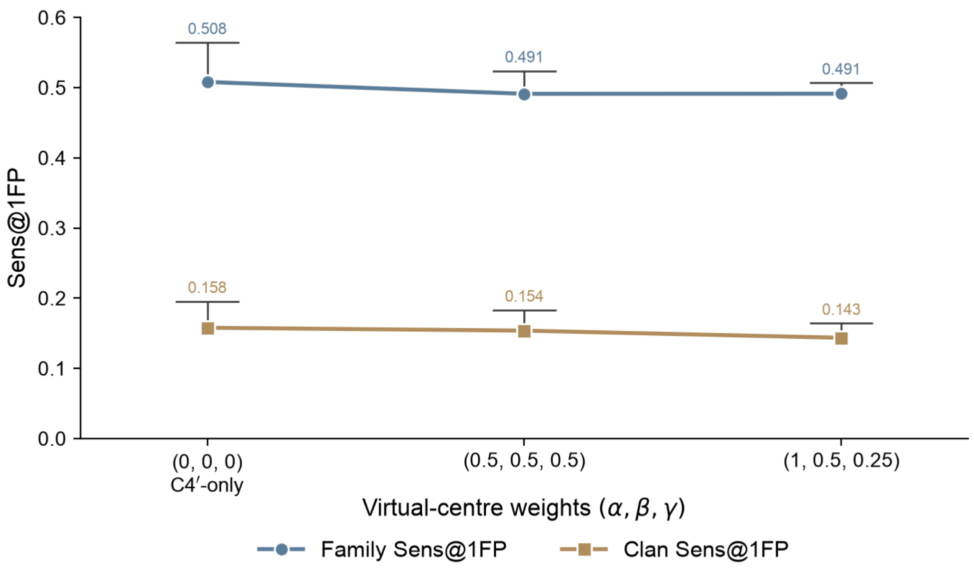


**Fig. S7**: **Virtual-centre weight ablation.**
Family-level and clan-level Sens@1FP for three virtual-centre weight settings $(\alpha, \beta, \gamma)$, evaluated by PDB-disjoint 5-fold cross-validation on the train–validation partition. All experiments use the fixed retrieval settings specified in Table S8. Points show the mean and error bars the standard deviation across the five folds.

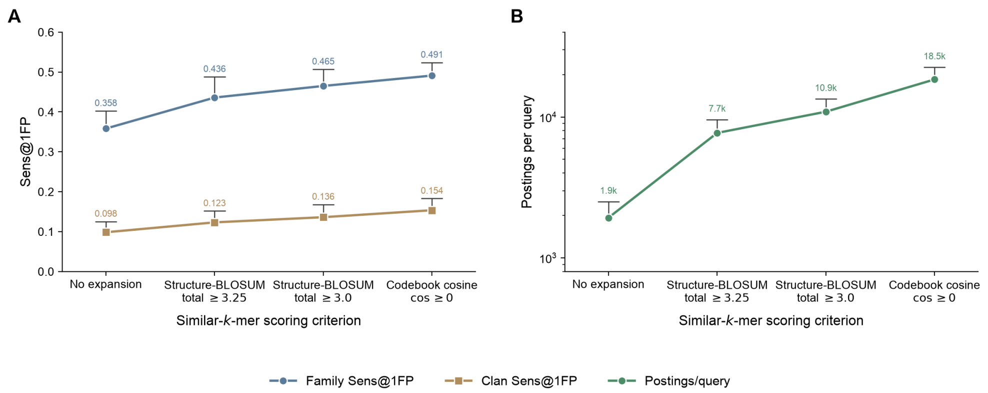


**Fig. S8**: **Similar-$k$-mer scoring trades prefilter sensitivity for selectivity.** **(A)** Family- and clan-level Sens@1FP for different criteria for admitting one-substitution neighbours of query 4-mers into the index lookup. Structure-BLOSUM criteria use the summed score over all four positions. **(B)** Mean postings traversed per query (log scale), counted before the double-diagonal filter and top-100 cutoff. All variants share the $K = 80$ VQ-VAE, contiguous 4-mer seed and fold-specific Structure-BLOSUM Smith–Waterman reranking. Mean ± s.d. over five PDB-disjoint folds.

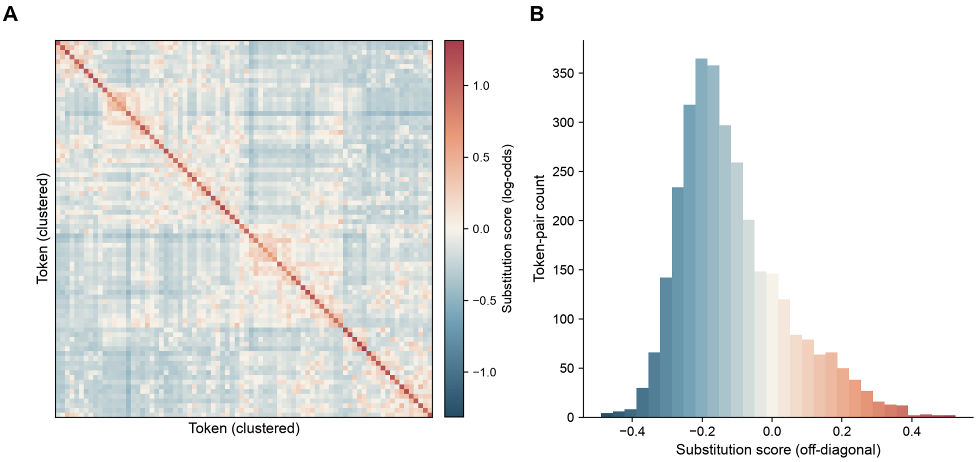


**Fig. S9**: **Structure-BLOSUM substitution matrix of RFoldSearch.**
**(A)** The $80 \times 80$ log-odds matrix used for similar-$k$-mer expansion and token Smith–Waterman scoring, estimated from aligned token pairs in training RNA pairs with symmetric US-align TM-score $\geq 0.5$ and scaled to a mean diagonal score of one. **(B)** Distribution of off-diagonal scores.

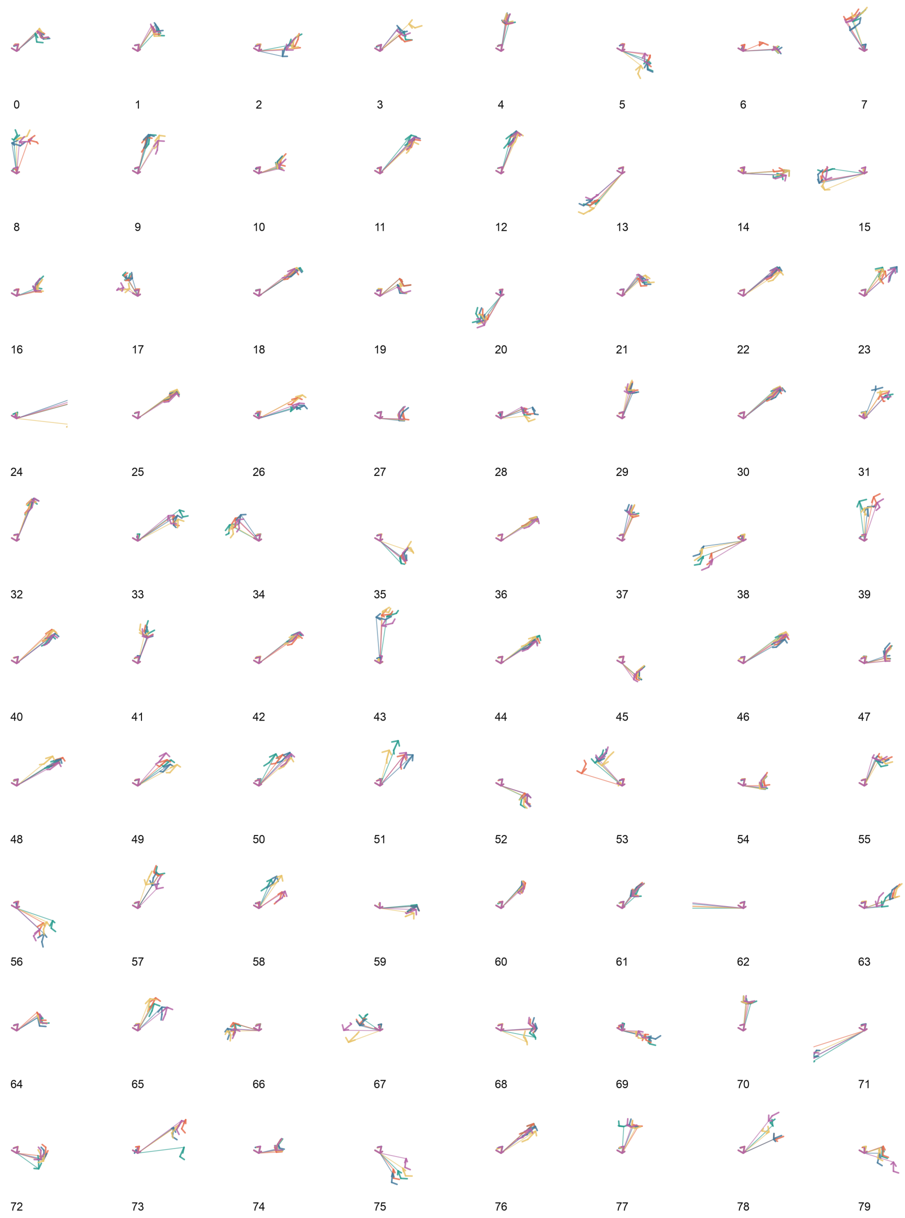


**Fig. S10**: **Visualisation of the 80 learned structural-alphabet states.** For each token, representative occurrences were selected by proximity to the empirical centroid of all assigned nucleotides in the normalised 17-dimensional RFoldSearch descriptor space. Each occurrence comprises a central nucleotide and its nearest nonlocal spatial neighbour. The pairs were aligned using the C3′–C4′–O4′ local frame of the central nucleotide, with C4′ as the origin. The five closest representatives, restricted to at most one occurrence per RNA chain, are overlaid for each state.

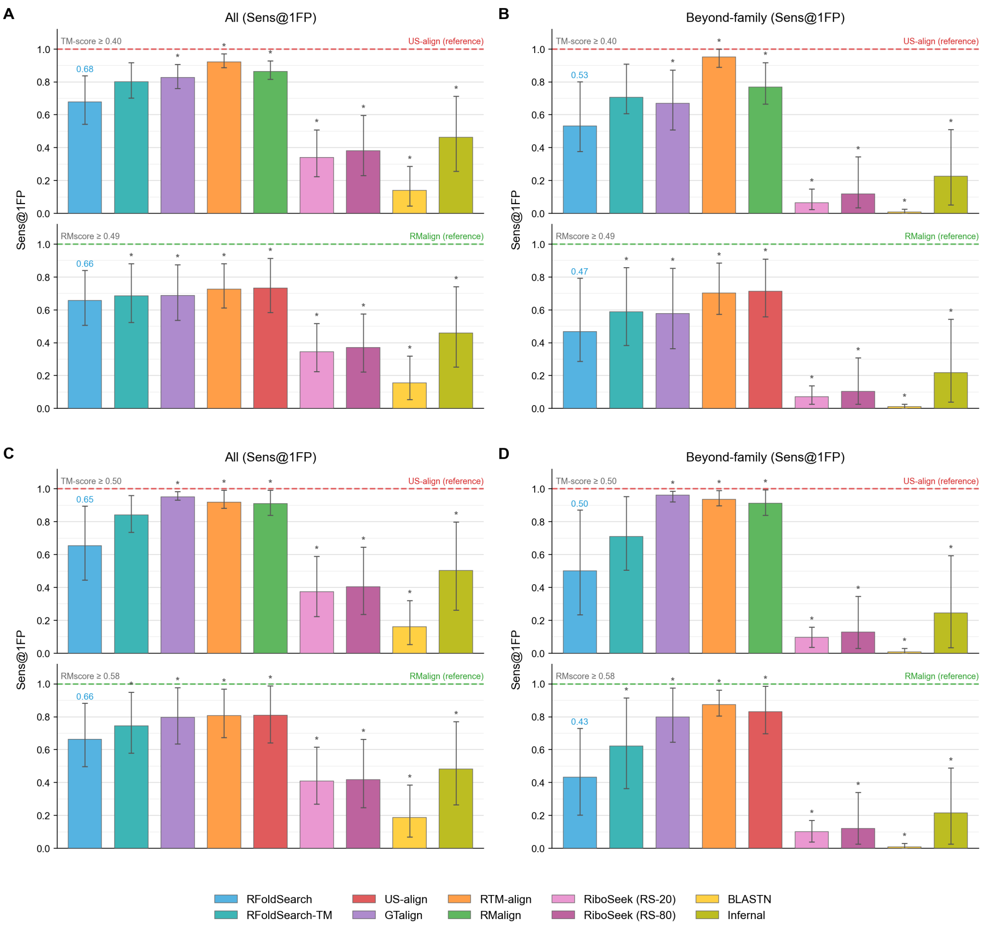


**Fig. S11**: **Structure-defined retrieval at alternative similarity thresholds.** Sens@1FP on the held-out RNAsolo 2.0 test set at symmetric US-align TM-score thresholds of (**A**, **B**) 0.40 and (**C**, **D**) 0.50, complementing Fig. 2. Upper and lower plots use TM-score- and RMscore-defined positives, respectively, with quantile-matched RMscore thresholds of 0.49 and 0.58. The corresponding reference aligner (US-align or RMalign) is shown as a dashed line rather than a bar. Bars indicate mean Sens@1FP with 95% block-bootstrap confidence intervals; asterisks indicate significant paired differences from RFoldSearch.

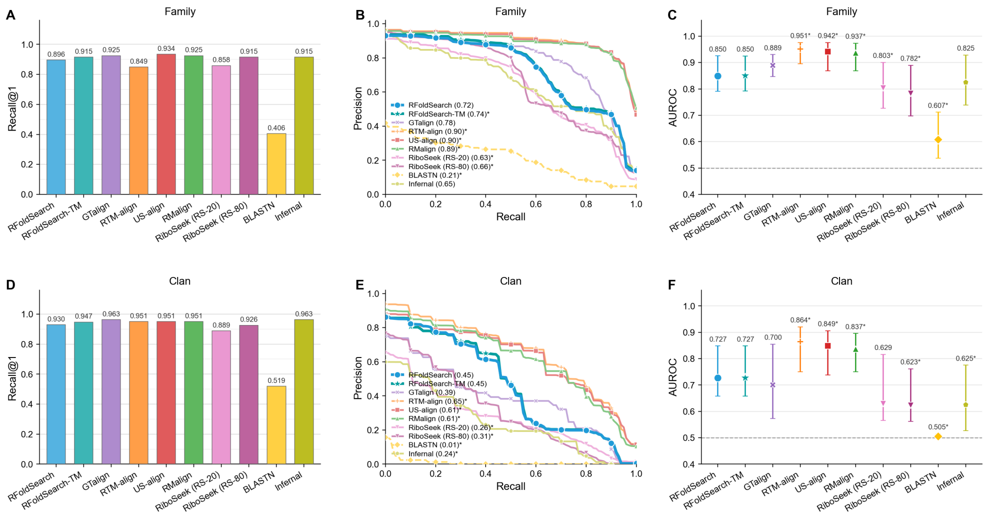


**Fig. S12**: **Supplementary retrieval metrics on the RNAsolo benchmark.** **(A–C)** Family-level and **(D–F)** clan-level retrieval performance on held-out queries. **(A,D)** Recall@1. **(B,E)** Query-averaged precision–recall curves, with MAP reported in the legend. **(C,F)** Mean per-query AUROC with 95% confidence intervals; the dashed line marks chance (0.5). Targets not returned by a method were treated as tied below all returned targets. Asterisks indicate significant paired differences in (B,E) or (C,F) relative to RFoldSearch. Confidence intervals and paired differences were obtained from 2,000 hierarchical block-bootstrap replicates resampling Rfam families or clans.

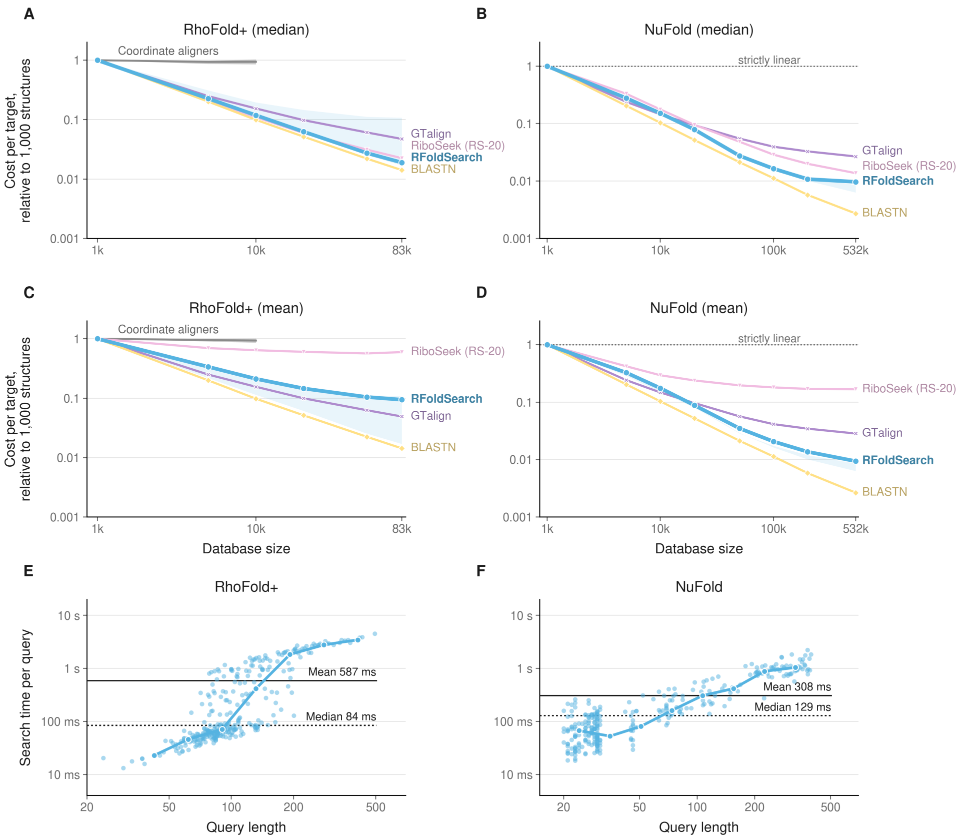


**Fig. S13**: **Runtime scaling with database size and query length.**
**(A–D)** Scaling of per-target search cost with database size on (A,C) RhoFold+ and (B,D) NuFold, calculated using median (A,B) or mean (C,D) query times. Per-target search cost was normalised to the corresponding value at 1,000 database structures. The horizontal dotted line at 1 represents strictly linear scaling of total query time, while values below 1 indicate lower per-target costs relative to the baseline. The grey band represents the range of coordinate aligners, and the shaded region around RFoldSearch indicates the interquartile range. RiboSeek used RS-20, and GTalign was run on a GPU. **(E,F)** Relationship between query length and RFoldSearch runtime on the full (E) RhoFold+ and (F) NuFold databases, with 300 queries each. Each point represents an individual query. Solid and dotted horizontal lines indicate mean and median runtimes, respectively, while blue trend lines connect binned medians.

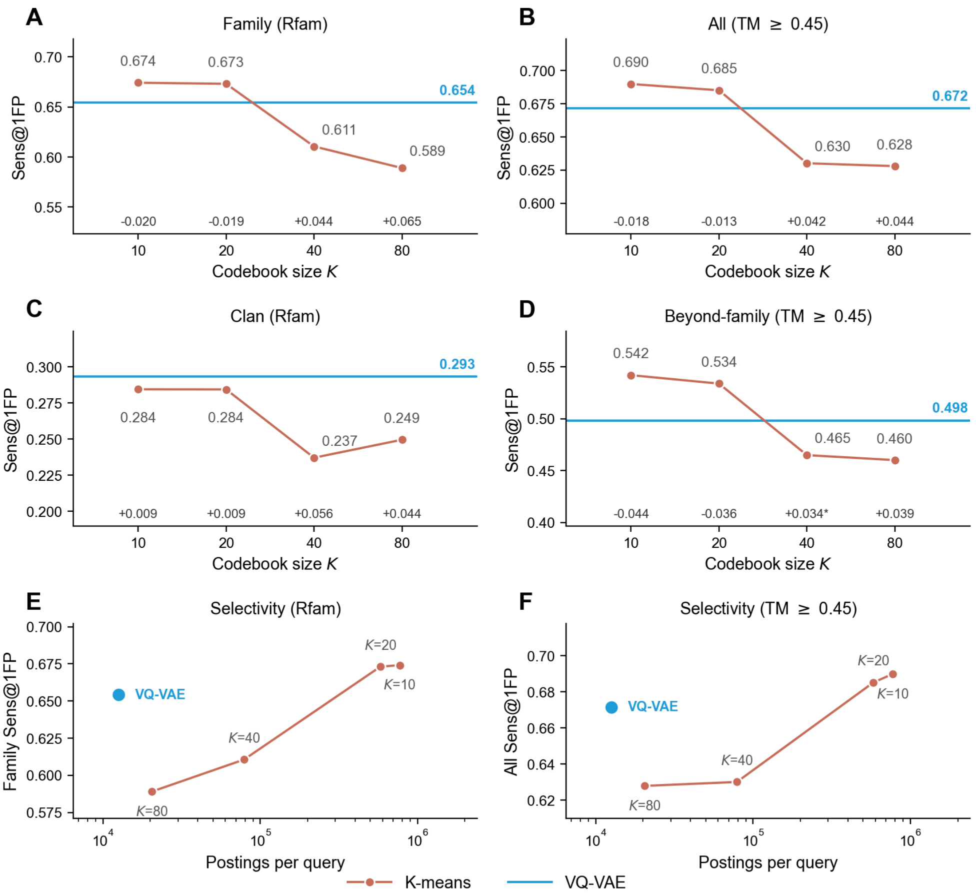


**Fig. S14**: **Quantisation control: $k$-means versus the learned VQ-VAE.** The same 17-dimensional descriptor was quantised by $k$-means or the VQ-VAE, and each alphabet was searched with the RFoldSearch pipeline and its own Structure-BLOSUM matrix. **(A, B)** Family-level and **(C, D)** clan-level Sens@1FP against codebook size. Horizontal lines mark the VQ-VAE; numbers along the bottom give paired differences (VQ-VAE minus $k$-means), with asterisks indicating 95% block-bootstrap intervals excluding zero. **(E, F)** Family-level Sens@1FP against prefilter selectivity, measured as the mean number of inverted-index postings traversed per query.

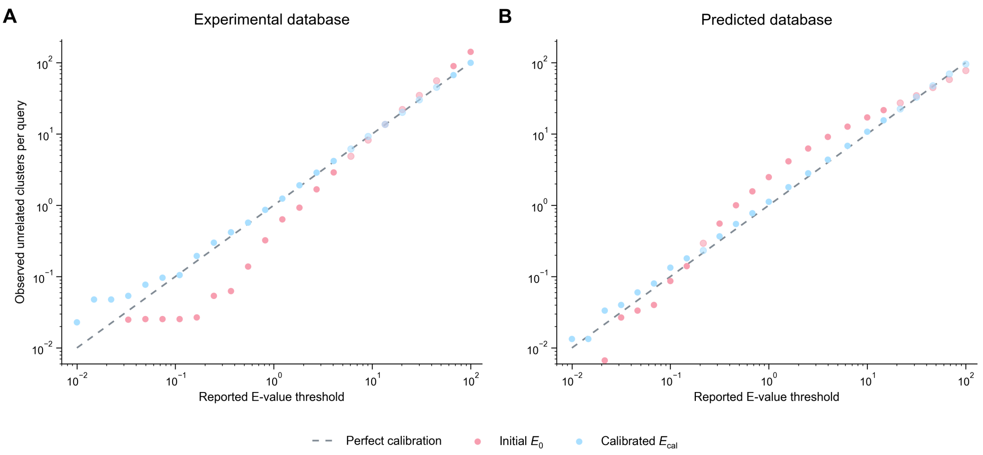


**Fig. S15**: **Calibration of RFoldSearch E-values.**
Calibration of RFoldSearch statistical significance on experimental and predicted RNA-structure databases. **(A)** Experimental database containing 885 RNA chains from RNAsolo 2.0, scored using raw token Smith–Waterman alignment. **(B)** Predicted database containing 33,043 RNA structures from NuFold, scored using window-20 composition-adjusted alignment. Each point plots the family-balanced mean number of retained background clusters per query against the corresponding reported E-value threshold. Pink points show the initial Gumbel E-value $E_0$, and light-blue points show the empirically calibrated value $E_{\mathrm{cal}}$. The dashed diagonal denotes perfect calibration, where the observed count equals the reported threshold.

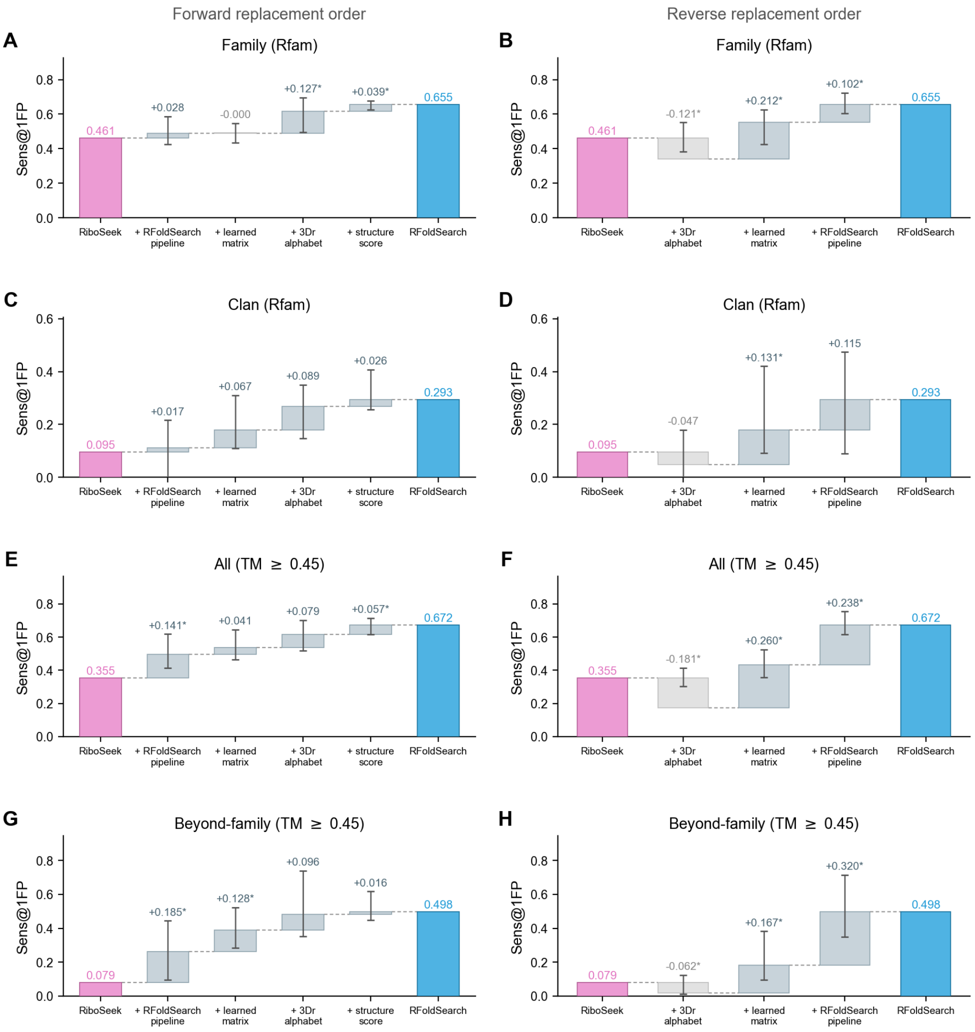


**Fig. S16**: **Attribution of the performance difference between RiboSeek and RFoldSearch.**
Starting from RiboSeek, components were replaced one at a time in (**A**, **C**, **E**, **G**) forward or (**B**, **D**, **F**, **H**) reverse order: the search pipeline, Structure-BLOSUM, the 3Dr alphabet and the structure score. Sens@1FP was evaluated at the (A, B) Rfam family and (C, D) clan levels and with structure-defined positives for (E, F) all and (G, H) beyond-family targets. Floating bars show each step; error bars are 95% paired block-bootstrap intervals over (A, B, E, F) Rfam families or (C, D, G, H) clans; asterisks mark intervals excluding zero.

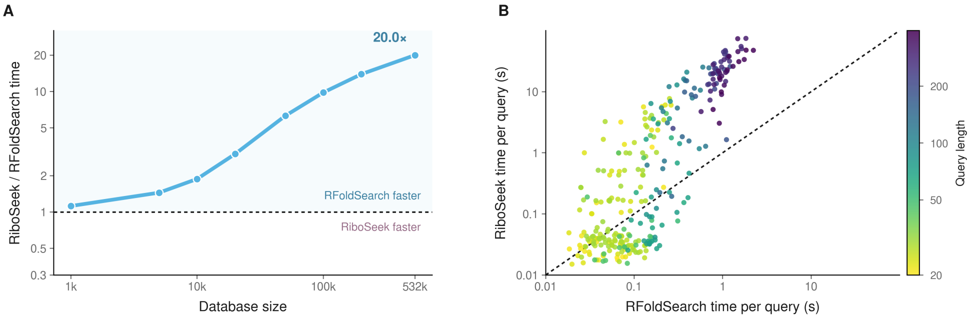


**Fig. S17**: **Runtime comparison of RFoldSearch and RiboSeek.** RiboSeek used RS-20, the alphabet used in the accuracy benchmarks. **(A)** Ratio of mean query runtimes (RiboSeek/RFoldSearch) across NuFold database sizes. The dotted line marks equal runtime, and shading indicates where RFoldSearch is faster. **(B)** Per-query runtime comparison on the full NuFold database, coloured by query length.

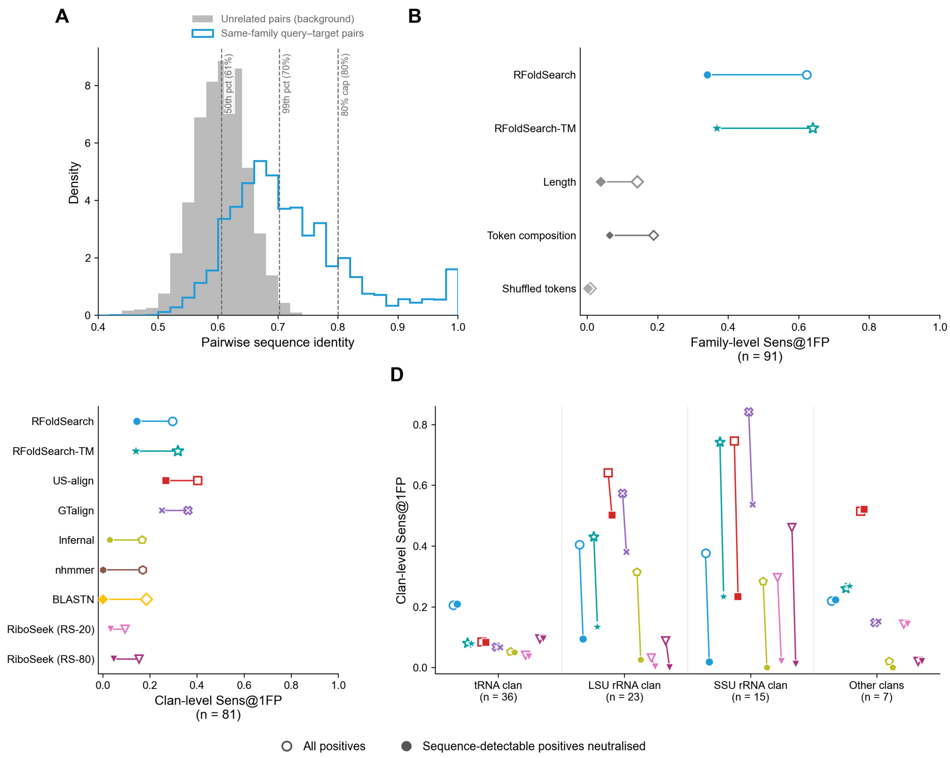


**Fig. S18**: **Controls and clan-level analysis of sequence-divergent retrieval.** **(A)** Sequence-identity distributions of 5,000 unrelated RNA pairs and of same-family query–target pairs. Dashed lines mark the background median (61%), 99th percentile (70%) and the 80% upper limit that define the identity ranges in Fig. 2E. **(B)** Family-level Sens@1FP of RFoldSearch, RFoldSearch-TM and negative controls that rank targets by length difference, 3Dr-token composition similarity or shuffled query tokens. **(C)** Clan-level Sens@1FP over all clan queries. **(D)** Clan-level Sens@1FP by clan. In B–D, open symbols use all positives and filled symbols make positives detected by BLASTN or nhmmer at $E \leq 10$ neutral.

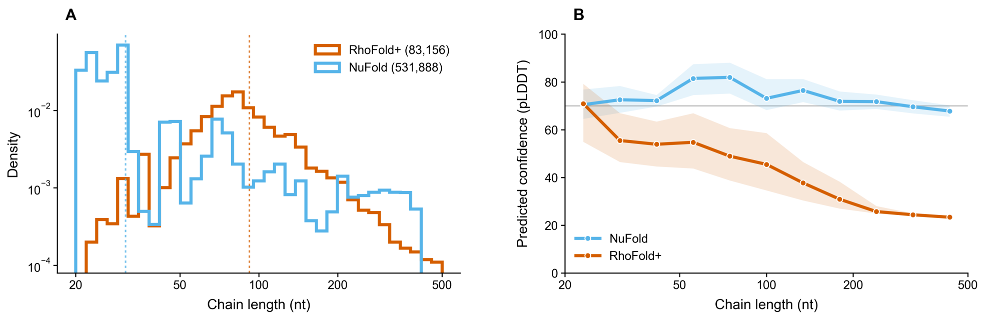


**Fig. S19**: **Composition and predicted confidence of the two predicted-structure databases.**
**(A)** Chain-length distributions of the RhoFold+ and NuFold databases. **(B)** Median pLDDT within logarithmic chain-length bins, with shaded interquartile ranges. RhoFold+ pLDDT values were multiplied by 100 to match the NuFold scale. The grey line indicates a pLDDT of 70.

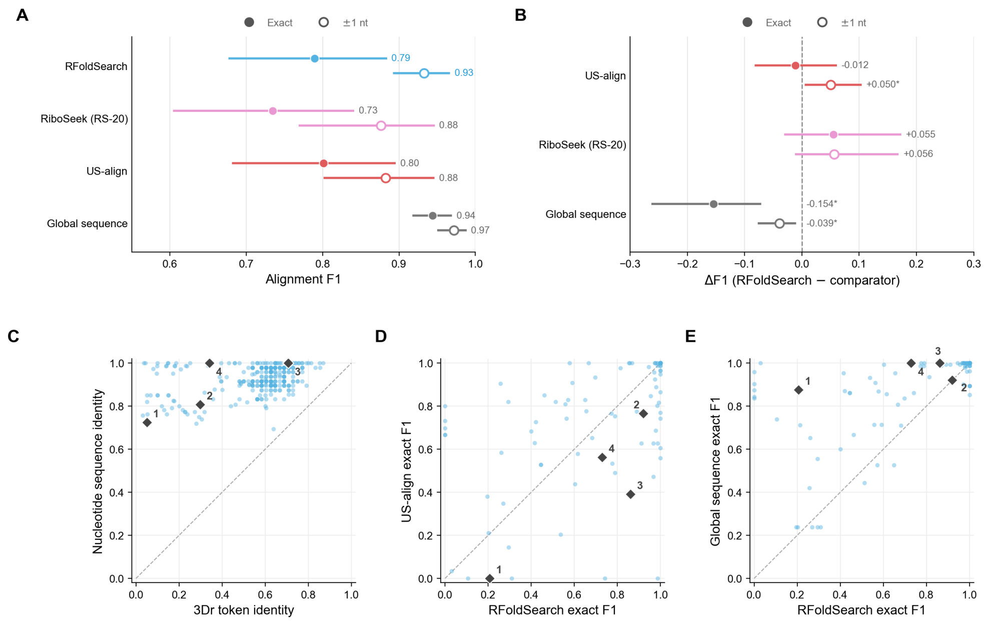


**Fig. S20**: **Nucleotide-level alignment accuracy against Rfam 3D seed alignments. (A)** Alignment $F_1$ for RFoldSearch, RiboSeek (RS-20), US-align and global sequence alignment. Filled and open markers indicate exact matching and ±1-nt tolerance, respectively. Points show family-macro means with 95% family-block bootstrap confidence intervals. **(B)** Paired differences in $F_1$, with asterisks indicating 95% confidence intervals excluding zero. **(C)** Nucleotide sequence identity versus 3Dr token identity. **(D,E)** Exact-match $F_1$ for RFoldSearch versus (D) US-align and (E) global sequence alignment. In C–E, each point represents a structure pair and dashed diagonals indicate numerical equality. Numbered diamonds identify four RF00015 pairs sharing 5O9Z chain 4 as the query, with corresponding alignment examples shown in Fig. S21.

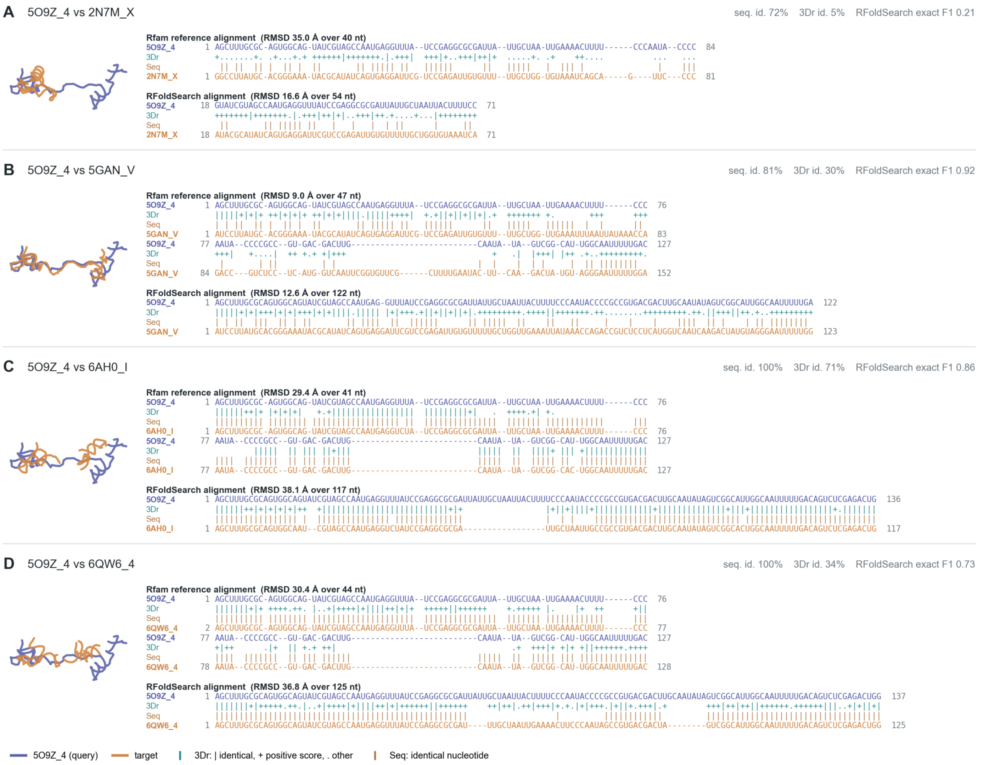


**Fig. S21: Examples of local structural alignments produced by RFoldSearch.**
Four pairs from Rfam family RF00015 are shown, all using 5O9Z chain 4 as the query. Panels **A**–**D** correspond to diamonds 1–4 in Fig. S20C–E. For each pair, the header reports nucleotide-sequence identity, 3Dr token identity and the exact $F_1$ agreement between the RFoldSearch alignment and the Rfam reference. The structure view on the left shows the query and target C4′ traces superposed according to the Rfam correspondence. On the right, the upper alignment is the Rfam 3D seed reference and the lower alignment is generated independently by aligning the two 3Dr token sequences with RFoldSearch. The reported RMSD is calculated from the C4′ atom pairs defined by each alignment. In the `3Dr` row, `|` indicates identical tokens, `+` indicates different tokens with a positive Structure-BLOSUM score, and `.` indicates token pairs with a non-positive score. In the `Seq` row, `|` indicates identical nucleotides.

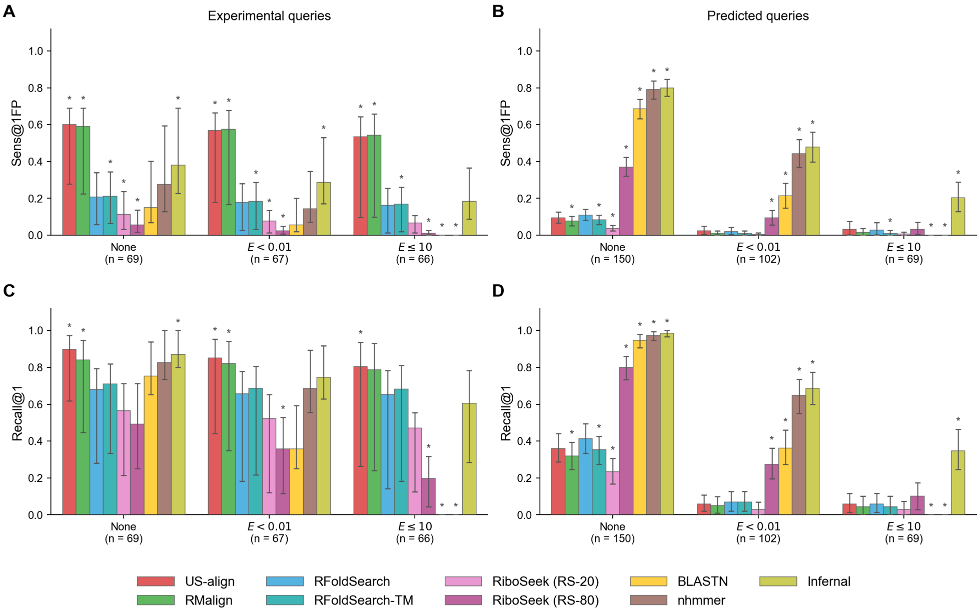


**Fig. S22**: **Retrieval from the predicted-structure database under Rfam-family ground truth.** (**A**,**B**) Sens@1FP and (**C**,**D**) Recall@1 for (**A**,**C**) experimental and (**B**,**D**) predicted queries. The x axis indicates whether all positives are retained (None) or positives detected by BLASTN or nhmmer at $E < 0.01$ or $E \leq 10$ are treated as neutral. Bars show query means, and error bars show 95% confidence intervals from 2,000 hierarchical bootstrap replicates resampling Rfam families. Asterisks mark methods whose paired difference from RFoldSearch has a 95% confidence interval excluding zero.

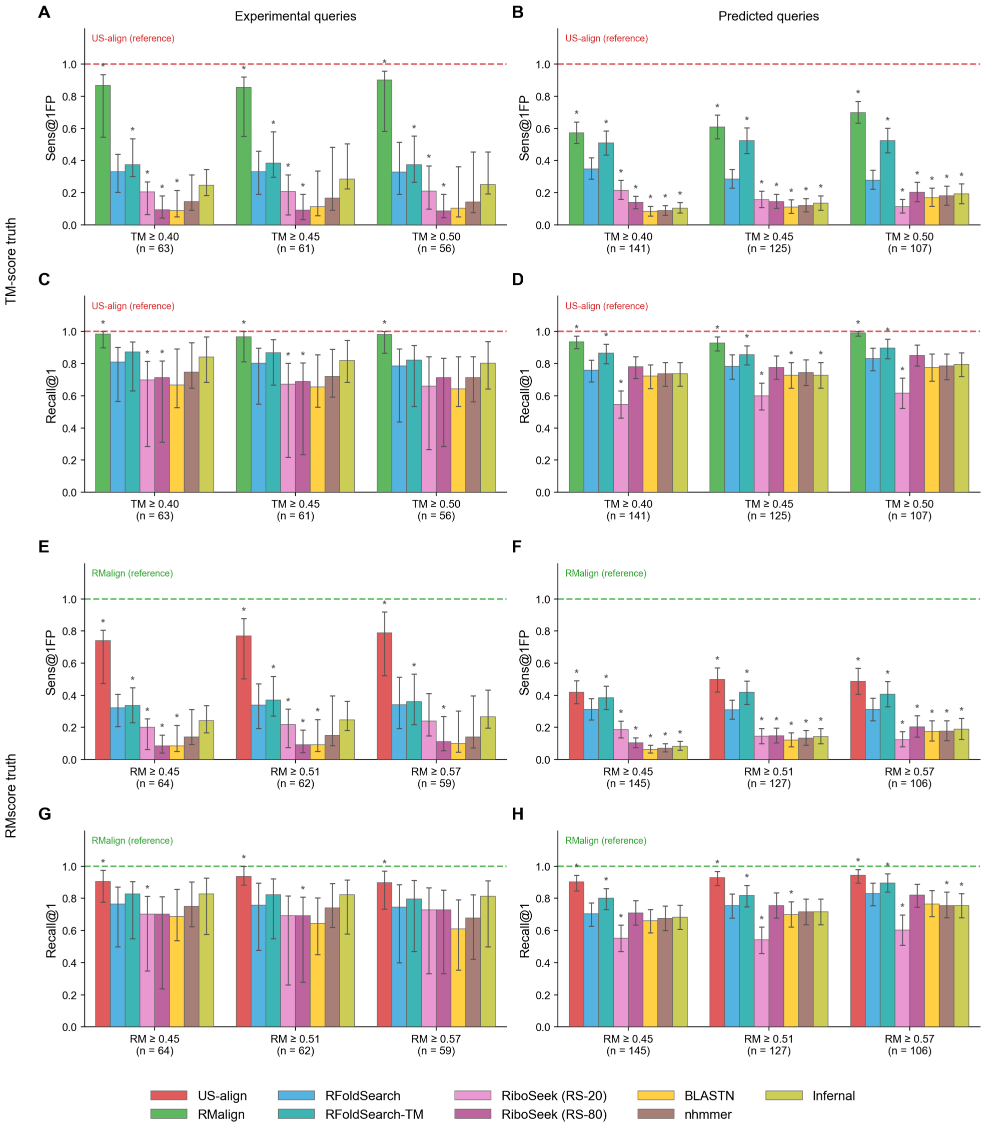


**Fig. S23**: **Retrieval from the predicted-structure database under structure-defined ground truths.** **(A–D)** TM-score truth. **(A,B)** Sens@1FP. **(C,D)** Recall@1. **(E–H)** RMscore truth. **(E,F)** Sens@1FP. **(G,H)** Recall@1. Under each truth, the aligner that defines the positives (US-align for TM-score, RMalign for RMscore) is shown as a dashed reference line at 1.00, and the other exhaustive aligner is evaluated as a competing method. Bars show query means, and error bars show 95% confidence intervals from 2,000 hierarchical bootstrap replicates resampling Rfam families. Asterisks mark methods whose paired difference from RFoldSearch has a 95% confidence interval excluding zero.

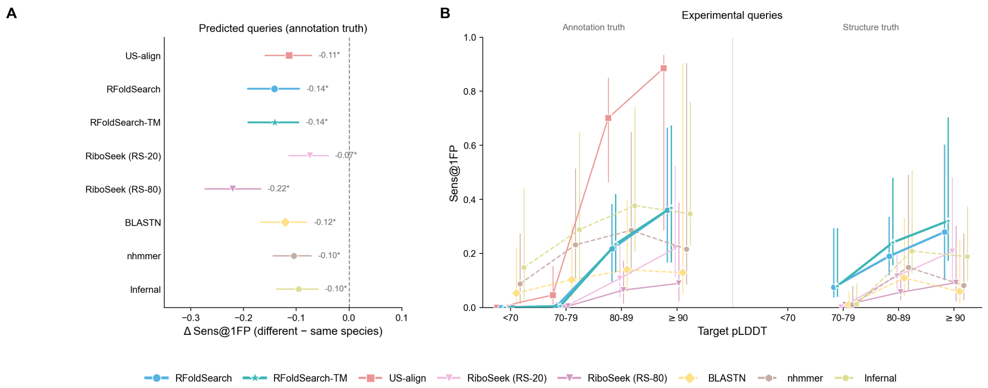


**Fig. S24**: **Retrieval from the predicted-structure database by organism and prediction confidence.** **(A)** Effect of cross-organism retrieval for predicted queries under annotation-defined truth. Asterisks mark differences whose 95% confidence interval excludes zero. **(B)** Sens@1FP of experimental queries, stratified by target pLDDT, under annotation-defined truth and structure-defined truth (symmetric US-align TM-score $\geq 0.45$). Error bars show 95% confidence intervals from 2,000 hierarchical bootstrap replicates resampling Rfam families.

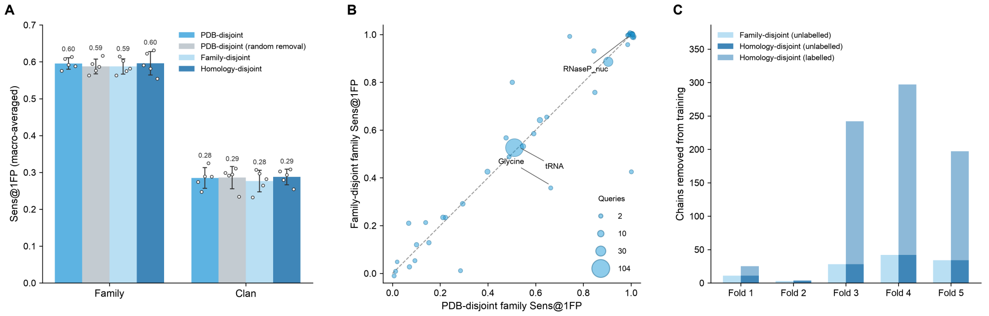


**Fig. S25**: **Family-disjoint and homology-disjoint cross-validation.**
**(A)** Family- and clan-level Sens@1FP under PDB-disjoint, random-removal, family-disjoint and homology-disjoint splits, macro-averaged over 41 families and 7 clans. Bars show means over five seeds, error bars ± s.d. and dots individual seeds. **(B)** Seed-averaged family-level Sens@1FP by family for PDB-disjoint and family-disjoint splits; marker area indicates query count. **(C)** Numbers of training chains excluded per fold in the family-disjoint and homology-disjoint schemes.

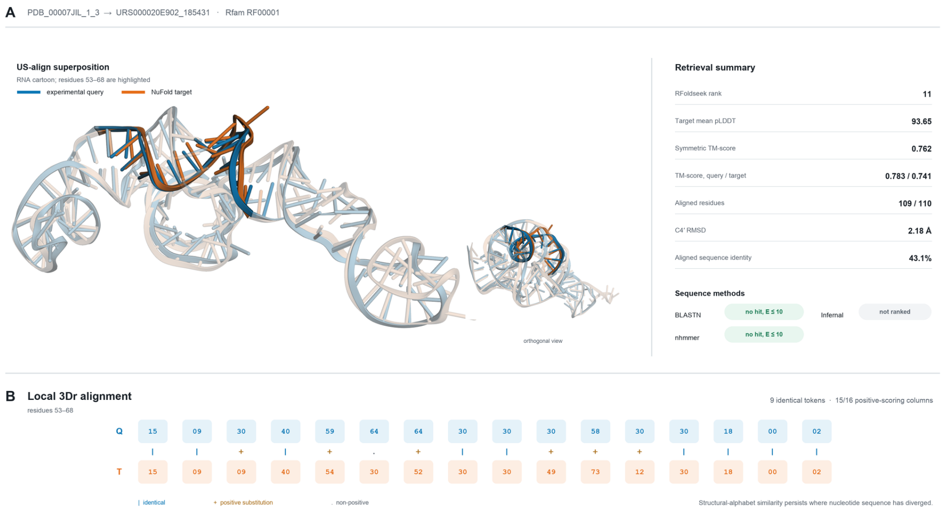


**Fig. S26**: **A sequence-invisible 5S rRNA homologue recovered by RFold-Search. (A)** US-align superposition of the experimental query `PDB_00007JIL_1_3` and the NuFold target `URS000020E902_185431`; nucleotides 53–68 are highlighted. The aligned nucleotide identity was 43.1%, whereas the symmetric TM-score was 0.762 over 109 aligned nucleotides. **(B)** The corresponding 3Dr-token alignment for nucleotides 53–68. Nine of 16 tokens were identical and 15 of 16 aligned token pairs had positive substitution scores.

#### Supplementary Tables

<table>
<tr><th>Partition</th><th>Chains</th><th>Rfam-labelled</th><th>Rfam families</th><th>Rfam clans</th><th>Mean length</th></tr>
<tr><td>Train</td><td>610</td><td>359</td><td>82</td><td>25</td><td rowspan="2">394$^\dagger$</td></tr>
<tr><td>Validation</td><td>85</td><td>43</td><td>17</td><td>7</td></tr>
<tr><td>Test</td><td>190</td><td>114</td><td>35</td><td>11</td><td>417</td></tr>
<tr><td>Total</td><td>885</td><td>516</td><td>96</td><td>25</td><td>—</td></tr>
</table>

**Table S1**: **Dataset and split summary.**
The corpus is partitioned at the PDB level into training, validation and held-out test sets at an approximate 80/20 test-holdout ratio (see "Methods – Data partitioning"). Training and validation are used jointly for tokeniser training and for 5-fold cross-validation over all model-design and ablation choices. *Rfam-labelled* counts chains carrying at least one Rfam family annotation; *Rfam families/clans* counts the distinct Rfam families/clans represented in each partition. $^\dagger$Pooled mean length across the 695 training+validation chains; per-partition values are not reported separately.

| Category | Parameter | Fixed setting |
|---|---|---|
| Data split | Target pool | Train + Validation |
| Data split | Query set | 5-fold cross-validation |
| Tokeniser | Method | $k$-means |
| Tokeniser | $k$-means settings | `batch_size`= 10,000, `max_iter`= 200, `n_init`= 3, `random_state`= 42 |
| Retrieval | Prefilter | Exact contiguous $k$-mer inverted index ($k = 4$) |
| Retrieval | Candidate set size | Top-100 |
| Retrieval | Reranking | Token Smith–Waterman alignment |
| Retrieval | Substitution matrix | Cluster-centroid cosine |
| Retrieval | Gap open / extend | $-2.0$ / $-0.5$ |
| Evaluation | Metrics | Family Sens@1FP, Clan Sens@1FP |

**Table S2: Fixed settings for the feature-group ablation experiments.**

| Category | Parameter | Setting |
|---|---|---|
| Data split | Target pool | Train + Validation |
| Data split | Query set | 5-fold cross-validation |
| Descriptor | Active blocks | $\mathbf{g}_{\mathrm{ctx}} + \mathbf{g}_{\mathrm{nn}}^{(1)}$ (17-D) |
| Descriptor | Virtual-centre weights $(\alpha, \beta, \gamma)$ | (0.5, 0.5, 0.5) |
| Tokeniser | Method | VQ-VAE |
| Model | Encoder / decoder depth | 2 / 4 |
| Model | Hidden / latent dimension | 128 / 128 |
| Model | Vector quantisation | EMA codebook update (decay 0.99) |
| Model | Commitment loss weight $\beta$ | 1.0 |
| Training | Batch size | 512 nucleotides |
| Training | Maximum epochs | 50 |
| Training | Optimiser | AdamW |
| Training | Learning rate | $3 \times 10^{-4}$ |
| Training | Weight decay | 0.01 |
| Training | Gradient clipping (L2 norm) | 1.0 |
| Training | Checkpoint selection | Lowest fold-internal validation loss |
| Retrieval | Prefilter | Similar-$k$-mer expansion inverted index, codebook-cosine neighbours |
| Retrieval | Candidate set size | Top-100 |
| Retrieval | Reranking | Token Smith–Waterman alignment |
| Retrieval | Substitution matrix | Codebook-cosine |
| Retrieval | Gap open / extend | −2.0 / −0.5 |
| Evaluation | Metrics | Family Sens@1FP, Clan Sens@1FP |

**Table S3**: **Fixed settings for the VQ-VAE codebook-size sweep.**

| Category | Parameter | Setting |
|---|---|---|
| Data split | Target pool | Train + Validation |
| Data split | Query set | 5-fold cross-validation |
| Descriptor | Active blocks | $\mathbf{g}_{\mathrm{ctx}} + \mathbf{g}_{\mathrm{nn}}^{(1)}$ (17-D) |
| Descriptor | Virtual-centre weights $(\alpha, \beta, \gamma)$ | (0.5, 0.5, 0.5) |
| Model | Codebook size $K$ | 80 |
| Model | Vector quantisation | EMA codebook update (decay 0.99) |
| Model | Commitment loss weight $\beta$ | 1.0 |
| Training | Batch size | 512 nucleotides |
| Training | Maximum epochs | 50 |
| Training | Optimiser | AdamW |
| Training | Learning rate | $3 \times 10^{-4}$ (cosine decay, 200 linear warm-up steps) |
| Training | Weight decay | 0.01 |
| Training | Gradient clipping (L2 norm) | 1.0 |
| Training | Checkpoint selection | Lowest fold-internal validation loss |
| Retrieval | Prefilter | Similar-$k$-mer expansion inverted index ($k = 4$), codebook-cosine neighbours |
| Retrieval | Candidate set size | Top-100 |
| Retrieval | Reranking | Token Smith–Waterman alignment |
| Retrieval | Substitution matrix | Codebook-cosine |
| Retrieval | Gap open / extend | $-2.0$ / $-0.5$ |
| Evaluation | Metrics | Family Sens@1FP, Clan Sens@1FP |

**Table S4**: **Fixed settings for the VQ-VAE architecture ablation.**

| Config. | $N_{\text{enc}}$ | $N_{\text{dec}}$ | $d_{\text{hidden}}$ | $d_{\text{latent}}$ |
|---|---|---|---|---|
| A1 | 1 | 3 | 128 | 128 |
| A2 | 1 | 3 | 256 | 128 |
| A3 | 2 | 2 | 128 | 128 |
| A4 | 2 | 4 | 128 | 128 |
| A5 | 2 | 8 | 128 | 128 |
| A6 | 2 | 8 | 256 | 128 |
| A7 | 4 | 8 | 256 | 128 |
| A8 | 4 | 8 | 256 | 64 |

**Table S5**: **Eight VQ-VAE architecture configurations tested in the ablation study.**

| Category | Parameter | Fixed setting |
|---|---|---|
| Data split | Target pool | Train + Validation |
| Data split | Query set | 5-fold cross-validation |
| Descriptor | Active blocks | $\mathbf{g}_{\mathrm{ctx}} + \mathbf{g}_{\mathrm{nn}}^{(1)}$ (17-D) |
| Tokeniser | Method | VQ-VAE |
| Model | Codebook size $K$ | 80 |
| Model | Encoder / decoder depth | 2 / 4 |
| Model | Hidden / latent dimension | 128 / 128 |
| Model | Vector quantisation | EMA codebook update (decay 0.99) |
| Model | Commitment loss weight $\beta$ | 1.0 |
| Training | Learning rate | $3 \times 10^{-4}$ (cosine decay, 200 linear warm-up steps) |
| Training | Weight decay | 0.01 |
| Training | Gradient clipping (L2 norm) | 1.0 |
| Training | Checkpoint selection | Lowest fold-internal validation loss |
| Retrieval | Prefilter | Similar-$k$-mer expansion inverted index ($k = 4$), codebook-cosine neighbours |
| Retrieval | Candidate set size | Top-100 |
| Retrieval | Reranking | Token Smith–Waterman alignment |
| Retrieval | Gap open / extend | $-2.0$ / $-0.5$ |
| Evaluation | Metrics | Family Sens@1FP, Clan Sens@1FP |

**Table S6**: **Fixed settings for the substitution-matrix ablation.**

| Category | Parameter | Fixed setting |
|---|---|---|
| Data split | Target pool | Train + Validation |
| Data split | Query set | 5-fold cross-validation |
| Descriptor | Active blocks | $\mathbf{g}_{\mathrm{ctx}} + \mathbf{g}_{\mathrm{nn}}^{(1)}$ (17-D) |
| Tokeniser | Method | VQ-VAE |
| Model | Codebook size $K$ | 80 |
| Model | Encoder / decoder depth | 2 / 4 |
| Model | Hidden / latent dimension | 128 / 128 |
| Model | Vector quantisation | EMA codebook update (decay 0.99) |
| Model | Commitment loss weight $\beta$ | 1.0 |
| Training | Checkpoint selection | Lowest fold-internal validation loss |
| Prefilter | Seed length ($k$, informative positions) | 4 |
| Prefilter | Diagonal double-match requirement | At least 2 hits per diagonal offset |
| Retrieval | Candidate set size $N$ | Top-100 |
| Retrieval | Reranking | Token Smith–Waterman alignment |
| Retrieval | Substitution matrix | Structure-BLOSUM diagmean-1 |
| Retrieval | Gap open / extend | $-2.0$ / $-0.5$ |
| Evaluation | Metrics | Family Sens@1FP, Clan Sens@1FP |

**Table S7**: **Fixed settings for the prefilter ablation.**
**E4** contiguous 1111 (baseline exact 4-mer, window $w = 4$); **G4** gapped 11_11 ($w = 5$); **S4** spaced 1_1_1_1 ($w = 7$); **SimE4** exact 4-mer with 1-substitution similar-$k$-mer expansion. All use $k = 4$ informative positions, the same reranker and the same top-100 candidates. Neighbours in SimE4 are admitted by codebook cosine similarity; the choice of neighbour criterion is examined separately in Fig. S8.

| Category | Parameter | Fixed setting |
|---|---|---|
| Data split | Target pool | Train + Validation |
| Data split | Query set | 5-fold cross-validation |
| Descriptor | Active blocks | $\mathbf{g}_{\mathrm{ctx}} + \mathbf{g}_{\mathrm{nn}}^{(1)}$ (17-D) |
| Tokeniser | Method | VQ-VAE |
| Model | Codebook size $K$ | 80 |
| Model | Encoder / decoder depth | 2 / 4 |
| Model | Hidden / latent dimension | 128 / 128 |
| Model | Vector quantisation | EMA codebook update (decay 0.99) |
| Model | Commitment loss weight $\beta$ | 1.0 |
| Training | Checkpoint selection | Lowest fold-internal validation loss |
| Retrieval | Prefilter | Similar-$k$-mer expansion ($k = 4$), codebook-cosine neighbours |
| Retrieval | Candidate set size | Top-100 |
| Retrieval | Reranking | Token Smith–Waterman alignment |
| Retrieval | Substitution matrix | Structure-BLOSUM diagmean-1 |
| Retrieval | Gap open / extend | $-2.0$ / $-0.5$ |
| Evaluation | Metrics | Family Sens@1FP, Clan Sens@1FP |

**Table S8: Fixed settings for the virtual-centre weight ablation.**

| Top_N | Sens@1FP | | Recall@1 | | AUPR | | Time |
|---|---|---|---|---|---|---|---|
| | family | clan | family | clan | family | clan | (rel.) |
| 100 | 0.545 | 0.194 | 0.872 | 0.931 | 0.653 | 0.426 | ×1.0 |
| 200 | 0.575 | 0.217 | 0.872 | 0.931 | 0.728 | 0.499 | ×1.4 |
| 300 | 0.574 | 0.206 | 0.872 | 0.931 | 0.776 | 0.513 | ×1.6 |
| 500 | 0.575 | 0.215 | 0.872 | 0.931 | 0.814 | 0.531 | ×1.9 |

**Table S9**: **RFoldSearch candidate-pool depth ablation.**
Effect of the candidate-pool depth on retrieval, measured on the validation split. Times are per-query search cost relative to top-$N$=100; they were measured with the reference implementation, so the ratios indicate how cost grows with $N$ for that implementation rather than for the released package, whose cheaper prefilter would make the relative increase steeper. Absolute per-query costs for the released package are in Figs. 3A and B.

| | RhoFold+ (83,156) | | NuFold (531,888) | |
|---|---|---|---|---|
| Method | Time (s) | Memory (GB) | Time (s) | Memory (GB) |
| RFoldSearch | 0.587 | 5.5 | 0.308 | 12.7 |
| RiboSeek (RS-20) | 7.081 | 94.6 | 6.172 | 97.5 |
| RiboSeek (RS-80) | 0.103 | 1.4 | 0.086 | 7.0 |

**Table S10**: **Search time and peak memory on the full predicted-structure databases.**
Mean per-query search time over 300 queries on one CPU core, and peak resident set size of one search process that loaded the database, built its index and ran all queries. RFoldSearch keeps database coordinates in memory for structure scoring, and RiboSeek stores token sequences and an index of spaced seeds.
